# Structures and molecular mechanisms of RAD54B in modulating homologous recombination

**DOI:** 10.64898/2026.03.22.713441

**Authors:** Pengtao Liang, Stephanie Tye, Johanna Ertl da Costa, Neelam Maharshi, Bilge Argunhan, Lucas Kuhlen, Megan Battley, Elizabeth A. McCormack, Wolf-Dietrich Heyer, Markus Löbrich, Xiaodong Zhang

## Abstract

Genome stability is essential for cellular viability yet constantly threatened by endogenous and exogenous DNA-damaging agents. Among these, DNA double-strand breaks (DSBs) are particularly harmful and in S/G2 phases are faithfully repaired through homologous recombination (HR), a high-fidelity pathway utilising homologous sequences in sister chromatin. The RAD51 recombinase forms nucleoprotein filaments on single-stranded DNA (ssDNA) to mediate homology search, strand invasion and subsequent D-loop formation that leads to DNA synthesis and repair. The efficiency of HR depends on precise regulation of RAD51 filament dynamics by accessory factors, including RAD54 and RAD54B, which belong to the SWI2/SNF2-family DNA translocases. While RAD54 is well-characterized, RAD54B’s molecular functions remain poorly understood. Here, we define RAD54B’s role in HR using cryo-electron microscopy, mutagenesis, biochemical and cellular assays. We show that RAD54B stabilizes RAD51-DNA filaments, inhibits RAD51 ATPase activity, and promotes strand invasion, D-loop formation and strand exchange. The N-terminal domain (NTD) alone supports filament stabilization and strand exchange, while the C-terminal ATPase domain is required for D-loop formation. Structural and biochemical analyses reveal three RAD51-interacting sites within the NTD and a unique domain (β-domain) that bridges RAD51 protomers and contacts donor dsDNA. This β-domain also regulates RAD54B’s ATPase activity and higher-order oligomer organization on dsDNA. Cellular assays reveal that the NTD RAD51-interacting sites as well as the β-domain are required for repairing camptothecin-induced DSBs by HR in human cells. Our findings uncover a modular architecture and mechanistic framework for RAD54B function in HR, highlighting its critical role in genome maintenance.

**Highlights:**

1. cryoEM structure of RAD54B in complex with RAD51-DNA complex
2. RAD54B uses three sites to interact with RAD51, including a previously unrecognised β-domain that bridges distal RAD51 protomers.
3. The β-domain plays multiple crucial roles including regulating filament stability, RAD54B ATPase activity and RAD54B higher order assembly on DNA.
4. RAD54B employs a modular mechanism, with the N-terminal region engaing and stabilising RAD51 filaments, capturing of the homologous strands, whereas the ATPase motor domainrequired for homology search and strand invasion.
5. RAD54B N-terminus and β-domain are essential for HR-mediated repair of camptothecin-induced breaks in human cells.

## Introduction

Maintaining genome stability is fundamental to cellular functions. However, cells are constantly challenged by exogenous agents such as chemical mutagens and ionizing radiation and endogenous cellular metabolites that can induce a variety of DNA lesions (Chatterjee and Walker 2017). Among these, DNA double-strand breaks (DSBs) represent one of the most deleterious forms of damage, as they can directly lead to the loss of essential genetic information. Homologous recombination (HR) is a key pathway that repairs DSBs with high fidelity and also plays a critical role in stabilizing stalled or collapsed replication forks (Wright, Shah, and Heyer 2018; Symington and Gautier 2011). Dysregulated HR can compromise genome integrity and drive tumorigenesis (Prakash et al. 2015; Jeggo, Pearl, and Carr 2016).

HR is initiated when nucleolytic resection generates 3′ single-stranded DNA (ssDNA) overhangs at the DSB site. These ssDNA ends are rapidly coated by replication protein A and subsequently displaced by the RecA-family recombinase RAD51, or its meiosis-specific counterpart DMC1, which form helical nucleoprotein assemblies known as presynaptic filaments. These filaments perform the essential task of homology search by interrogating double-stranded DNA (dsDNA) for sequence complementarity. Upon locating a homologous sequence, the filament mediates strand invasion, pairing the ssDNA with its complementary strand in the dsDNA to form a displacement loop (D-loop), which then leads to DNA synthesis and subsequent repair (Wright, Shah, and Heyer 2018; Ranjha, Howard, and Cejka 2018). RAD51 filaments are therefore central components of the HR pathway.

The efficiency and accuracy of HR rely on precise regulation of RAD51-DNA filament assembly, dynamics, and disassembly by a range of accessory factors (Carver and Zhang 2021). Furthermore, the intrinsic recombinase activity of RAD51 is low and requires modulators to stimulate. Among these modulators, the SWI2/SNF2-family DNA translocases RAD54 and RAD54B are ATP-driven motor proteins that directly bind RAD51 or DMC1 filaments and stimulate recombination activities (Sarai et al. 2008; Sanchez et al. 2013; Tanaka et al. 2002; Crickard, Moevus, et al. 2020; Shin et al. 2025; Ceballos and Heyer 2011). While RAD54 has been extensively characterized as a DNA motor that stimulates strand invasion and D-loop formation, the molecular functions of RAD54B remain comparatively obscure. RAD54B shares ∼50% sequence identity with RAD54 and retains a conserved C-terminal ATPase motor domain but displays differences in N-terminal domain (NTD). Previous studies have shown that human RAD54B directly interacts with both RAD51 and DMC1 and can stimulate recombinase-driven strand invasion or strand exchange *in vitro* (Sarai et al. 2006; Wesoly et al. 2006). Very recently, RAD54B was shown to function in synthesis-dependent strand annealing, one of the two sub-HR pathways (Chan et al. 2026). Studies of budding yeast ortholog Rdh54 (also known as Tid1) showed that it plays a role in both mitotic and meiotic recombination (Chi et al. 2009; Santa Maria et al. 2013). In mitotic recombination Rdh54 antagonized Rad54 during Rad51-mediated recombination (Shah et al. 2020) and acts as an overall negative regulator of D-loop formation (Piazza et al. 2019). Rdh54 binds directly to Rad51 filaments and uses ATP hydrolysis to translocate along DNA, remodel recombinase–DNA complexes, and in some contexts, displace Rad51 (Chi et al. 2006).

Despite these insights, significant gaps remain in our understanding of RAD54B’s mechanism in HR. While its interaction with RAD51 and DNA is established, the structural basis of these interactions and their functional consequences during the various stages of HR are not well understood.

To address these knowledge gaps, we set out to characterize the molecular mechanisms and functions of RAD54B in HR, with a particular focus on its ability to modulate RAD51-ssDNA filaments, promote strand invasion, and facilitate strand exchange. Using cryo-electron microscopy (cryo-EM) combined with targeted mutagenesis, biochemical reconstitution and cellular assays, we define the molecular basis by which RAD54B engages and modulates RAD51 filaments to regulate HR. Our results show that RAD54B inhibits the ATPase activity of DNA-bound RAD51, stabilizes recombinase filaments, and promotes strand invasion, D-loop formation, and strand exchange, thereby facilitating efficient HR.

Specifically, we show that the RAD54B-NTD is sufficient to stabilize RAD51 filaments and promote strand exchange, but not D-loop formation, which requires the C-terminal ATPase domain. This domain-specific functionality suggests a modular architecture in which the NTD mediates key protein-protein and protein-DNA interactions, while the ATPase domain provides the motor activity necessary for DNA remodelling during strand invasion. Structural and mutational analyses further reveal that the NTD contains three distinct RAD51-interacting sites, which are essential for filament stabilization. Importantly, the third binding site (β-domain) that bridges two distal RAD51 protomers in the filament and interacts with dsDNA via a positively charged loop. We further reveal that β-domain can bind to donor dsDNA and promote strand invasion and D-loop formation. Further, we show that the β-domain regulates RAD54B DNA-dependent ATPase activity and promotes functional higher order organisation, suggesting cross talk between NTD and CTD. We also assess DSB repair by HR in human cells and show that RAD54B knock-out (KO) cells exhibit a defect in repairing camptothecin-induced DSBs. The repair defect is rescued after transfection with wild-type RAD54B but not with RAD54B containing deletions in the NTD RAD51-interaction domains or mutations in the β-domain. Together we demonstrate the three sites have distinct functions in regulating RAD51 and HR activities, providing a structural framework for understanding its role in HR.

## Results

### RAD54B enhances RAD51 recombination activities via direct interactions

Previous biochemical studies have shown that human RAD54B interacts with RAD51 (Tanaka et al. 2000). More recently, single-molecule analysis demonstrated that the yeast ortholog Rdh54 binds directly to RAD51-ssDNA filaments (Crickard, Kwon, et al. 2020). To investigate how RAD54B engages with RAD51, we carried out *in vitro* studies using purified proteins (Fig. S1A). We first assessed the interactions between RAD54B and RAD51, as well as its respective DNA filaments. As expected, RAD51 can be pulled down by RAD54B (Fig. 1A) and these interactions extended to both ssDNA and dsDNA filament forms, as evidenced by the altered migration patterns of the filaments during electrophoresis (Figs. 1B-C), indicative of complex formation. To visualize these complexes, we employed negative-stain electron microscopy. 2D class averages of images of RAD51-ssDNA or -dsDNA complexes alone showed regular helical filaments (Fig. 1D). Upon addition of RAD54B, both types of filaments were decorated with additional density on the side of the filaments (Fig. 1E). Closer inspection of the 2D class averages revealed two-lobed densities (Fig. 1E, arrows), consistent with binding by RAD54B, which is predicted to have a bi-lobed structure consisting of two RecA-like domains. Interestingly, contrary to previous studies suggesting that RAD54B preferentially associates with filament ends (Sarai et al. 2006), our data suggest that RAD54B can bind along the length of RAD51 filaments.

**Figure 1.**
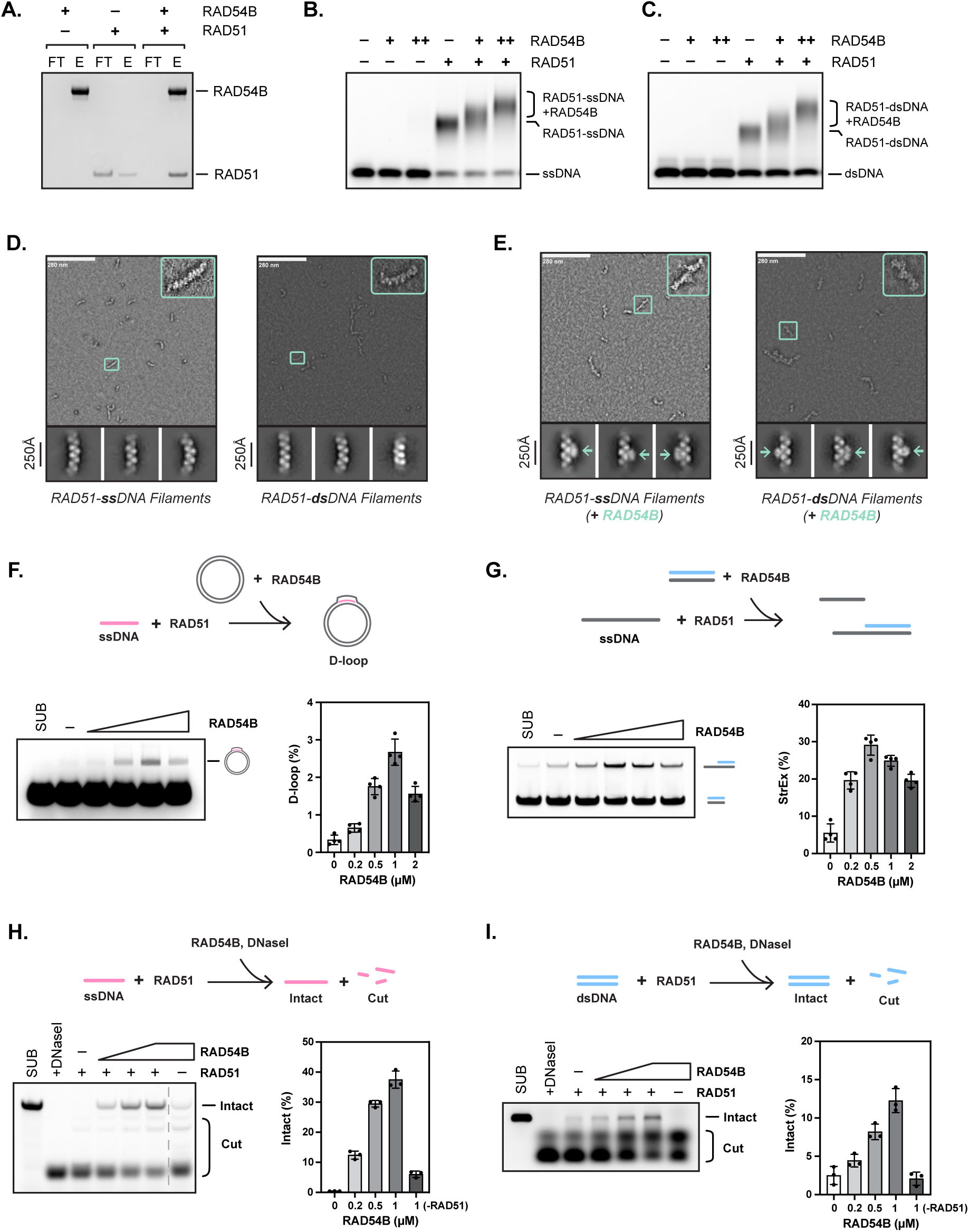
RAD54B stimulates RAD51-mediated recombination by binding to its filament form. **(A)** Pull-down experiment of full-length RAD54B with RAD51, 1µM protein for each. Elution (E) and flow-through (FT). **(B)** Electrophoresis mobility shift assay (EMSA) showing RAD54B (0.5 and 1 µM) binding to RAD51-ssDNA filaments. **(C)** Electrophoresis mobility shift assay (EMSA) showing RAD54B (0.5 and 1 µM) binding to RAD51-dsDNA filaments. **(D)** Negative-stain electron microscope micrographs and 2D class averages of RAD51 -ssDNA or -dsDNA filaments. **(E)** Negative-stain electron microscope micrographs and 2D class averages of RAD51 -ssDNA or dsDNA filaments in the presence of full-length RAD54B. **(F)** D-loop assay. Schematic (top), representative gels (left), quantification (right) showing stimulation of RAD51-driven strand invasion (1 µM, 6 µMnt, 60 µMbp) by full-length RAD54B (0.2, 0.5, 1, and 2 µM). **(G)** Oligonucleotide strand exchange assay. Schematic (top), representative gels (left), quantification (right) showing stimulation of RAD51-driven strand exchange (1 µM, 3 µMnt, 1 µMbp) by full-length RAD54B (0.2, 0.5, 1, and 2 µM). **(H)** ssDNA Nuclease protection assay. Schematic (top), representative gels (left), quantification (right) showing RAD51-ssDNA (1 µM, 10 µMnt) filaments stabilisation by full-length RAD54B (0.2, 0.5, and 1 µM). **(I)** dsDNA Nuclease protection assay. Schematic (top), representative gels (left), quantification (right) showing RAD51-dsDNA (0.5 µM, 5 µMbp) filaments stabilisation by full-length RAD54B (0.2, 0.5, and 1 µM). For **(F-G)**, averages are shown with error bars depicting standard deviation (n = 4). For **(H-I)**, averages are shown with error bars depicting standard deviation (n = 3).

The yeast Rdh54 was shown to stimulate RAD51-driven strand invasion (Petukhova, Sung, and Klein 2000). To investigate if human RAD54B has similar effects, we used a plasmid DNA-based D-loop assay (Liu, Sneeden, and Heyer 2011). RAD51 was pre-incubated with a 90-nt fluorophore-labelled ssDNA designed to target a unique homologous sequence in the plasmid DNA. Upon successful strand invasion, the RAD51–ssDNA filament displaces the non-complementary strand of the plasmid, forming a D-loop. The fluorophore label allowed detection of stable D-loop products by electrophoresis. Under the conditions employed here, RAD51 alone produced no detectable stable D-loops (Fig. 1F). The addition of RAD54B enhanced D-loop formation in a concentration-dependent manner with peak enhancement at ∼1 µM.

To assess strand exchange activities upon strand invasion, we employed an oligonucleotide strand exchange assay. Here, RAD51–ssDNA filaments were first formed on 116-nt single-stranded DNA, and then incubated with homologous, fluorescently labelled 60-bp double-stranded DNA (dsDNA). During the reaction, the ssDNA pairs with the fluorescently-labelled complementary strand, displacing the non-complementary strand and generating a heteroduplex composed of 60-bp dsDNA and a 56-nt ssDNA overhang (Fig. 1G). Substrates and products can be resolved by electrophoresis due to their distinct sizes, as visualised via the fluorescent label. Consistent with the strand invasion results, RAD54B enhanced the RAD51-driven strand exchange in a concentration-dependent manner that peaks around 0.5-1 µM (Fig. 1G).

The stimulatory effects could be due to enhanced filament stabilisation as proposed for other RAD51 modulators such as RAD51AP1 and RAD51 paralogs (Amunugama, Groden, and Fishel 2013; Kuhlen et al. 2025b; Greenhough et al. 2025). We next investigated its effects on RAD51 filaments stability using nuclease protection assays. The results showed an increased stabilisation with increasing RAD54B concentrations (Figs. 1H-I).

### N-terminal domain of RAD54B is sufficient to modulate RAD51 filaments

RAD54B contains a largely disordered and highly varied N-terminal domain and a conserved C-terminal ATPase domain (Figs. S1B-C). Previous studies identified the NTD of human RAD54B to be responsible for mediating its interaction with RAD51 (Tanaka et al. 2000). To assess their functions in RAD51 recombination activities, we purified the NTD (residues 1–285) and the CTD (residues 286–910) separately. Consistent with previous reports, the NTD (RAD54B-N) efficiently pulled down RAD51, similarly to full length proteins (Figs. 2A). By contrast, CTD (RAD54B-C) showed only minimal interactions, similarly to the background signal.

**Figure 2.**
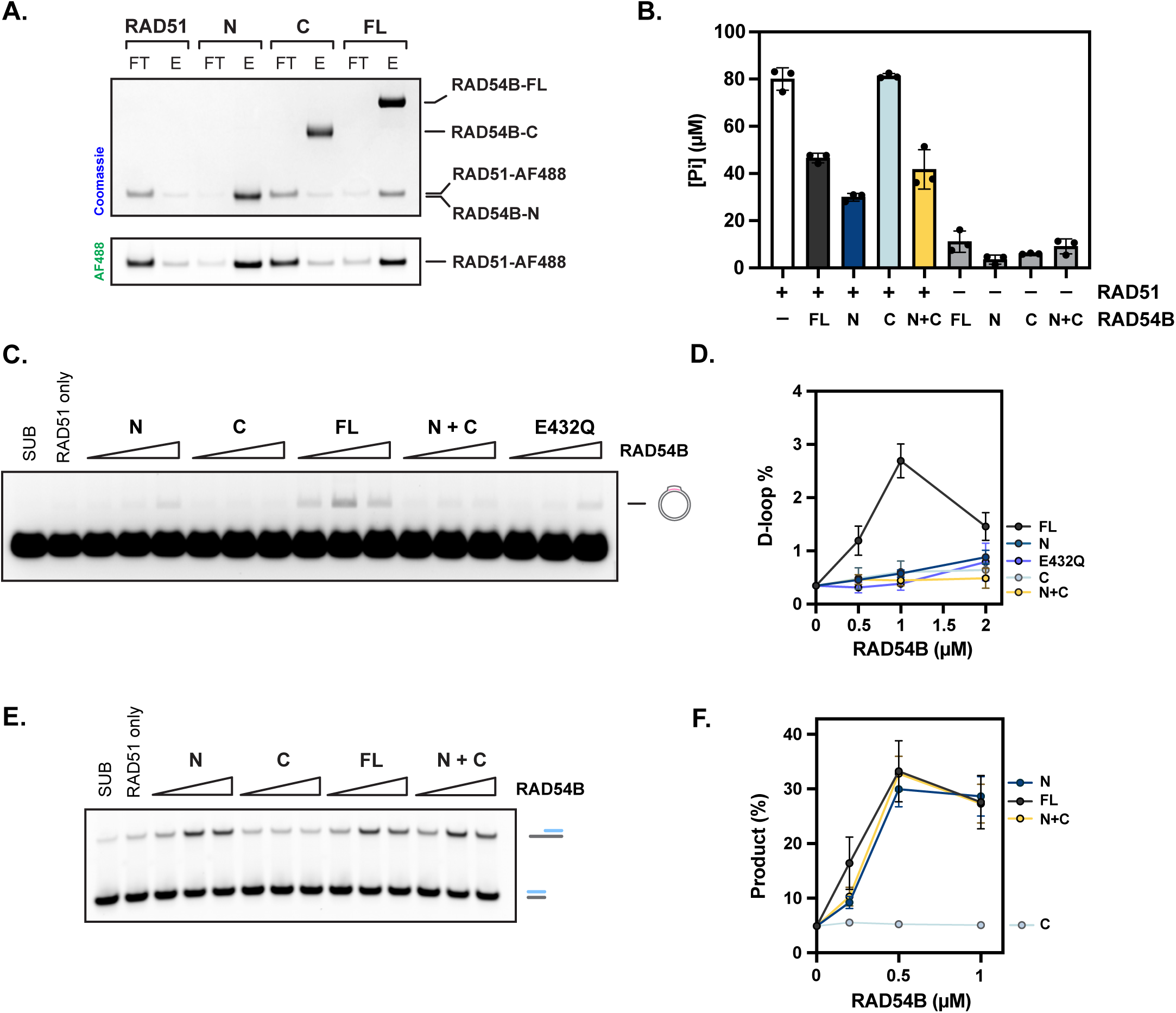
The N-terminal region of RAD54B is sufficient to recapitulate most activities in regulating RAD51. **(A)** Pull-down assay of RAD54B fragments (N-terminal, C-terminal, and full-length [FL]) with RAD51. Top gel was imaged after staining with Coomassie blue, bottom gel was imaged with Aurora Flour 488. **(B)** ATPase assay showing inhibition of RAD51 ATPase activity in ssDNA filaments by the N-terminal of RAD54B. **(C)** Representative gel images showing the stimulation of RAD51-driven D-loop formation by RAD54B (full-length, fragments, and mutant) (0.5, 1, and 2 µM). **(D)** Quantification of **(C)**. **(E)** Representative gel images showing the stimulation of RAD51-driven strand exchange by RAD54B (N, C, and FL) (0.2, 0.5, and 1 µM). **(F)** Quantification of **(E)**. For **(B-F)**, averages are shown with error bars depicting standard deviation (n = 3).

Some RAD51-interacting proteins regulate RAD51 ATPase activity; therefore, we examined whether RAD54B, and in particular its N-terminal domain (NTD), modulates RAD51 ATPase function. Both full-length RAD54B and RAD54B-N inhibited RAD51 ATPase activity when RAD51 was assembled into active filaments, whereas the C-terminal domain of RAD54B had no effect, consistent with its inability to bind RAD51 (Fig. 2B).

Since RAD54B-N recapitulates the binding and ATPase-modulating activity of the full-length protein, we asked whether it could also modulate RAD51-driven strand invasion, D-loop formation and strand exchange. In D-loop assays, both RAD54B-N and RAD54B-C failed to substitute for full-length RAD54B in promoting RAD51-mediated strand invasion (Figs. 2C–D). Notably, in oligonucleotide strand exchange assays, RAD54B-N recapitulated the activity of the full-length protein, whereas the C-terminal domain of RAD54B had no detectable effect (Figs. 2E–F). These results suggest that, while the NTD of RAD54B is able to efficiently promote strand exchange, the C-terminal ATPase domain is indispensable for promoting a stable D-loop formation. This notion is further supported by analysis of a RAD54B Walker B mutant (E432Q) that abolishes ATP hydrolysis. While this mutant retained the ability to stimulate strand exchange (Figs. S1D), it was defective in promoting D-loop formation (Figs. 2C-D, S2E-F), consistent with previous findings for RAD54 (Alexiadis, Lusser, and Kadonaga 2004) and highlighting the differences in these two assays in probing recombinase activities.

Taken together, our data demonstrate that RAD54B-N is essential and sufficient to mediate binding to RAD51, inhibit RAD51 ATPase activity and modulate strand exchange activities. However, full-length RAD54B, including an intact ATPase site, is required to support the complete functional activity of RAD54B in homologous recombination.

### Structural basis of RAD54B interaction with RAD51-DNA filaments

To provide a structural and molecular basis for how RAD54B engages and modulates RAD51–DNA filaments, we employed cryo-electron microscopy (cryo-EM) to visualize the complex between RAD54B-N and RAD51–DNA filaments.

We reconstituted RAD51-ssDNA filaments with RAD54B-N for cryo-EM analysis. Image processing produced 3D reconstructions of RAD51–ssDNA–RAD54B-N complexes at a global resolution of 2.6 Å in the presence of ATP hydrolysis-inhibited Ca^2+^-ATP (Figs. S2A) and 2.9 Å in the presence of Mg^2+^-ATP (Fig. S2B, 1^st^ Workflow Path). The two reconstructions are similar and have sufficient resolution that allowed us to build a structural model. Both reconstructions revealed densities corresponding to binding sites 1 and 2 of the RAD54B-N (Fig. 3A), contacting each RAD51 protomer (Fig. 3B, S3A). Site 1 involves the extreme N-terminal 10 residues of RAD54B including the Met1 which is acetylated in our proteins as confirmed by mass spectrometry. There are extensive and specific interactions: the acetylated methionine (AME1) inserts into a hydrophobic pocket on RAD51 while Arg2 forms a salt bridge with RAD51 residue Glu98. Additional hydrophobic interactions are mediated by Ala5, Ala6, and Leu10 from RAD54B and Phe20 and Ile148 from RAD51 (Fig. 3C). Site 2 includes an FxPP motif (^20^FIPP^23^) that is conserved in several RAD51 modulators including BRCA2, FIGNL1 and RAD51AP1 (Figs. 3A, 3D-E) (Miron et al. 2024), where Phe20 and Pro23 from RAD54B interact with a hydrophobic pocket on RAD51 filament surface (Fig. 3D).

**Figure 3.**
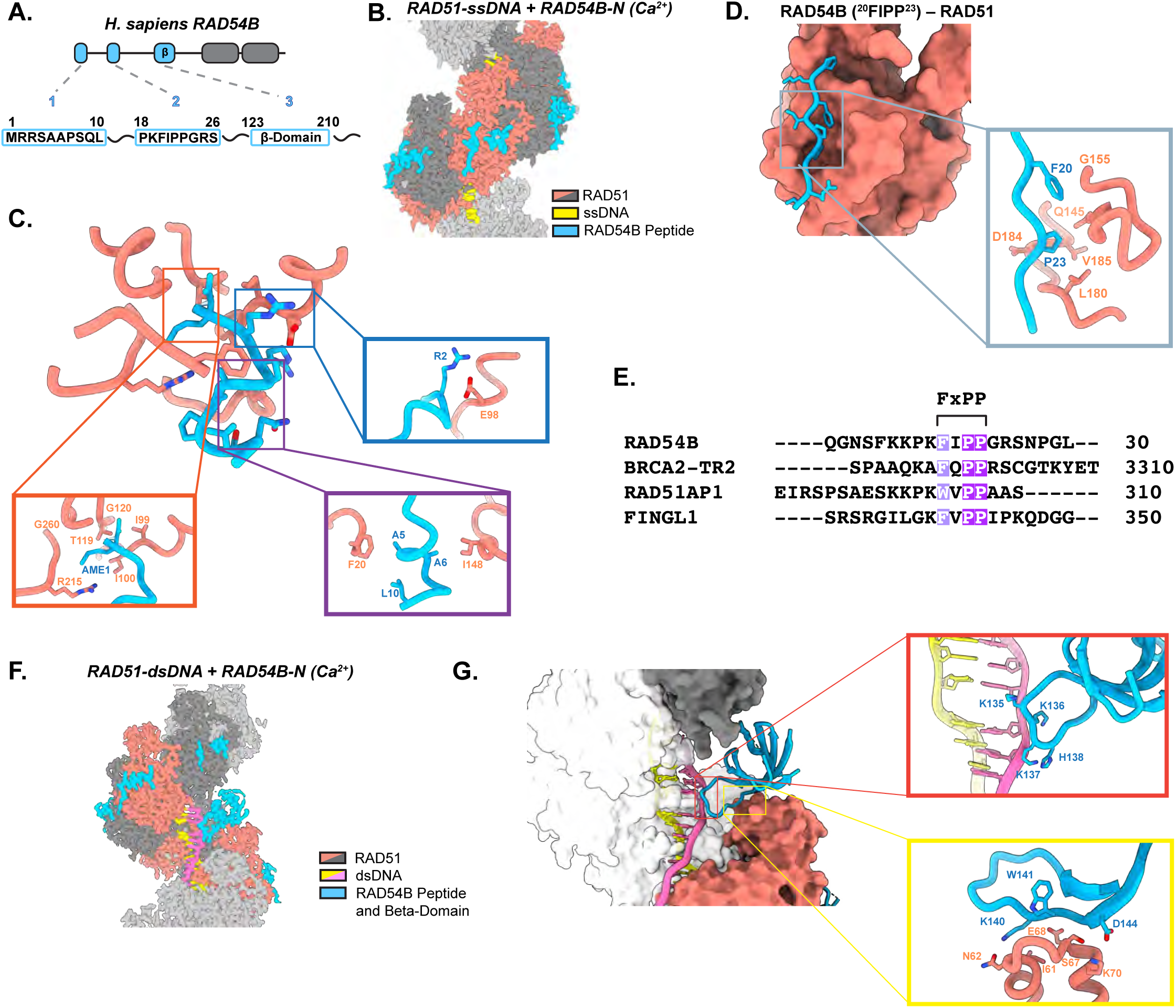
Cryo-EM structure of the RAD51-DNA filaments in complex with N-terminal domain of RAD54B. **(A)** Schematic of three distinct RAD51 binding sites in RAD54B-N. **(B)** Cryo-EM map of the Ca^2+^-ATP bound RAD51-ssDNA filament in the presence of RAD54B-N. **(C)** Closeup view of the RAD54B site 1 (blue) interacting with a RAD51 protomer (salmon). **(D)** Interaction between RAD54B site 2 (^20^FIPP^23^) (blue) and a RAD51 protomer (salmon). **(E)** Sequence alignment of RAD51 regulators highlighting the conserved FxPP motif. **(F)** Cryo-EM map of the Ca^2+^-ATP bound RAD51-dsDNA filament in the presence of RAD54B-N. **(G)** Structural details of the RAD54B β-domain interacting with RAD51-dsDNA filament.

In the Mg^2+^-ATP dataset, we obtained another 3D reconstruction, which contains a distinct globular density bridging two distal RAD51 protomers in the filaments (n and n+5) (Figs. S2B, 2^nd^ Workflow Path; S3B). Interestingly, the globular density was consistent with the AF3-predicted β-domain of RAD54B (Figs. S1C) (Abramson et al. 2024b). It inserts into the groove in the filaments and induces filament compression (Fig. S3C). However, due to the limited resolution in this region, detailed interaction features could not be discerned. The β-domain is located close to the ssDNA but there were no direct interactions that could be observed (Fig. S3B). We wondered whether it might contact dsDNA in the RAD51-dsDNA filaments. We obtained a 2.4 Å cryo-EM reconstruction of the RAD51–dsDNA–RAD54B-N complex in the presence of Ca²⁺-ATP (Fig. S4A-B). As with the ssDNA complex, RAD54B-N binds each RAD51 protomer via the same two sites in the extreme N-terminus (Fig. 3F, S3D). Notably, the β-domain was now clearly resolved bridging two distal RAD51 protomers (Fig. 3F). The AF3 model fitted into the density convincingly with density for the loop that contacts dsDNA clearly resolved (Fig. S3E). The β-domain contacts phosphor backbones of one of the two DNA strands via a positively charged loop (Fig. 3G), therefore resulting in a better resolved structure. In this structural model, there are weak electrostatic interactions between the β-domain and the two RAD51 protomers in the filament which would stabilise and compress the filaments.

Together, these cryo-EM structures reveal how RAD54B-N interacts with RAD51 filaments through three conserved motifs and suggest that the β-domain contributes to filament stabilization and dsDNA binding, potentially providing a molecular mechanism for the enhanced strand invasion activity conferred by RAD54B.

### The three binding sites in RAD54B have distinct functionalities

To corroborate our structural observations and assess the functional relevance of individual RAD54B–RAD51 interfaces, we used mutagenesis and biochemical assays to dissect the contributions of the three RAD51-binding sites. We generated three RAD54B deletion mutants: RAD54B-Δ10 (removal of residues 2–10, disrupting site 1 while retaining Met1), RAD54B-Δ25 (removal of residues 2–25, disrupting sites 1 and 2), and RAD54B-Δβ (removal of the β-domain, site 3) (Fig. 3A).

To understand how these deletions affected RAD51 binding. Compared with full-length RAD54B, deletion of site 1 (Δ10) nearly abolished binding to RAD51, whereas further removal of site 2 (Δ25) did not result in an additional detectable reduction (Fig. 4A-B). In contrast, deletion of the β-domain (Δβ) had no effect on RAD51 binding under these conditions, suggesting that this interaction is transient or of lower affinity. Together, these results indicate that RAD54B establishes its initial and dominant contact with RAD51 through site 1, which anchors RAD54B on the filament and positions the FxPP motif (site 2) and β-domain (site 3) for subsequent engagement.

**Figure 4.**
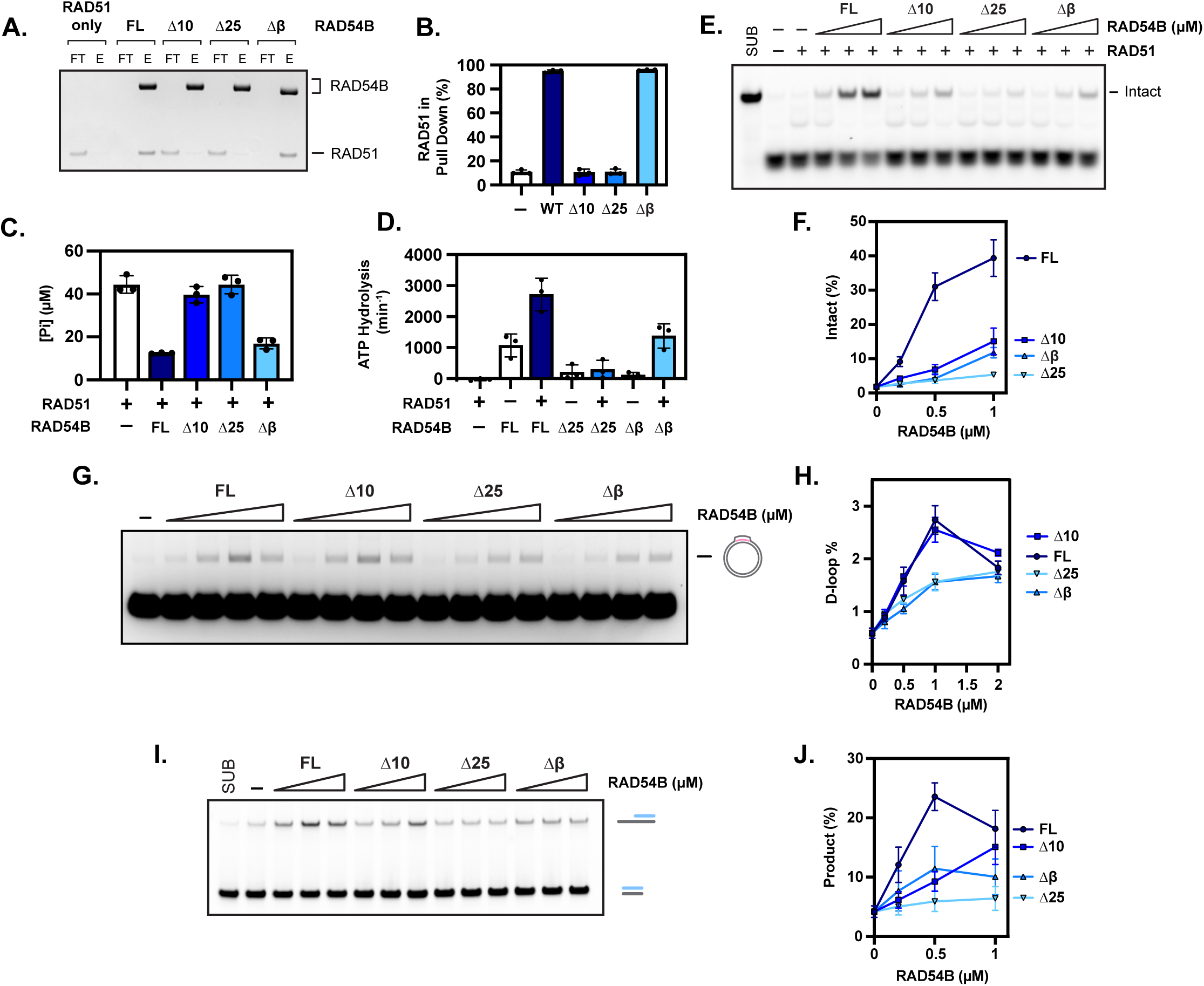
Three RAD51 binding sites in RAD54B serve distinct but complementary functions. **(A)** Pull-down experiment of RAD54B (full-length and deletion mutants) and RAD51. **(B)** Quantification of **(A)**. **(C)** ATPase assay showing that RAD54B inhibits RAD51 ATPase activity, compared across full-length (FL), Δ10, Δ25, and Δβ variants. **(D)** ATPase assay showing that RAD51 stimulates RAD54B-dsDNA dependent ATPase activity, compared across full-length (FL), Δ25, and Δβ variants. **(E)** ssDNA nuclease protection assay. Representative gel images showing RAD51-ssDNA filament stabilisation by RAD54B (full-length and deletion mutants). **(F)** Quantification of **(E)**. **(G)** Representative gel images showing the stimulation of RAD51-driven D-loop formation by RAD54B (full-length and deletion mutants) (0.2, 0.5, 1, and 2 µM). **(H)** Quantification of **(G)**. **(I)** Representative gel images showing the stimulation of RAD51-driven strand exchange by RAD54B (full-length and deletion mutants) (0.2, 0.5, and 1 µM). **(J)** Quantification of **(I)**. For **(A-J)**, averages are shown with error bars depicting standard deviation (n = 3).

We next assessed the impact of these mutations on RAD51 ATPase activity using RAD51-ssDNA filaments. Deletion of site 1 eliminated the ability of RAD54B to inhibit RAD51 ATP hydrolysis, whereas further deletion of site 2 had little additional effect. Removal of the β-domain did not affect RAD54B-mediated inhibition of RAD51 ATPase activity (Fig. 4C), indicating that site 1 is the principal determinant of ATPase suppression. Conversely, RAD51 stimulated the ATPase activity of RAD54B when RAD54B was bound to dsDNA (Fig. 4D). This stimulation was lost upon deletion of sites 1 and 2, consistent with their requirement for RAD51 binding. Notably, although deletion of the β-domain substantially reduced the intrinsic ATPase activity of RAD54B, the remaining activity could still be stimulated by RAD51, consistent with RAD51-mediated activation occurring through direct interaction with the N-terminal region of RAD54B.

We then examined the consequences of these mutations for filament stabilization. Deletion of site 1 (Δ10) markedly impaired stabilization of RAD51–ssDNA filaments (Fig. 4E-F). Structurally, site 1 inserts into the RAD51 N-terminal domain at the protomer–protomer interface, providing a direct mechanism for stabilizing filament architecture. In addition, site 1 is positioned adjacent to the ATP-binding pocket, offering an explanation for its inhibitory effect on ATP hydrolysis and its contribution to filament stabilization. This mechanism is consistent with the increased filament stability observed when ATP hydrolysis is suppressed by substitution of Mg²⁺ with Ca²⁺ to form non-hydrolysable Ca²⁺–ATP complexes (Spirek et al. 2018). As expected, deletion of both site 1 and site 2 (Δ25) eliminated filament stabilization (Fig. 4E-F). Notably, although deletion of the β-domain had minimal effects on RAD51 binding and ATPase inhibition, it severely compromised filament stabilization, consistent with a structural role in compressing RAD51 filaments and bridging adjacent RAD51 protomers.

We next evaluated how these mutations affected RAD51-mediated recombination. Despite its key roles in ATPase inhibition and filament stabilization, site 1 was dispensable for stimulating stable D-loop formation: full-length RAD54B and RAD54B-Δ10 supported comparable levels of D-loop formation (Fig. 4G-H). In contrast, deletion of either the first 25 residues (Δ25) or the β-domain (Δβ) substantially reduced D-loop stimulation. We then examined strand exchange using oligonucleotide-based assays. In contrast to the D-loop reaction, deletion of site 1 significantly reduced strand exchange stimulation and shifted the dose–response to higher RAD54B concentrations (Fig. 4I-J), consistent with impaired binding. Deletion of both site 1 and site 2 nearly abolished strand exchange stimulation, while deletion of the β-domain also reduced maximal activity.

Together, these results indicate that filament stabilization mediated by site 1 through inhibition of RAD51 ATPase activity plays distinct roles in strand invasion and strand exchange. RAD51-driven D-loop formation relies primarily on the RAD54B motor domain to translocate along dsDNA and promote strand invasion, rather than on suppression of RAD51 ATP hydrolysis. In contrast, efficient strand exchange requires inhibition of RAD51 ATPase activity, indicating that ATPase suppression is indispensable at this later stage of recombination. The FxPP motif (site 2) and the β-domain (site 3) appear to engage RAD51 filaments through transient interactions and promote filament stabilization by strengthening RAD51 protomer–protomer contacts, thereby contributing preferentially to strand invasion when the motor domain remains intact. Overall, these data indicate that RAD54B integrates binding, ATPase regulation, filament architecture, and motor activity through distinct structural elements to control discrete steps of homologous recombination.

### The β-domain of RAD54B plays key roles in homologous recombination via dsDNA binding and bridging distal RAD51 protomers

Structural analysis revealed that the β-domain of RAD54B bridges two distal RAD51 protomers within the filament, resulting in filament stabilization and compression. Notably, a positively charged loop within the β-domain (β-loop) was well resolved in the RAD51–dsDNA complex but was absent from the ssDNA-bound structure (Figs. S3E, 5A). Sequence alignment showed that this loop is highly conserved across eukaryotic species, highlighting its potential functional importance (Fig. S5A). Together, these observations suggested that the β-domain stabilizes RAD51–DNA filaments by bridging RAD51 protomers from adjacent helical turns and, in the context of dsDNA filaments, may additionally engage DNA via the β-loop (Fig. S5B).

To test this idea, the isolated β-domain was purified and examined in crosslinking assays with RAD51 in the presence of different DNA substrates. Upon mixing the β-domain with RAD51 or RAD51–DNA complexes, several new crosslinked species were detected (Figs. 5B, S5C). A prominent band of approximately 90 kDa, consistent with a complex containing two RAD51 protomers and a single β-domain, was markedly enriched in the presence of dsDNA compared with ssDNA or no DNA (Figs. 5B–C). To assess whether formation of this complex depends on β-loop–DNA interactions, four charged residues within the β-loop were mutated to alanine (β-domain-4A). This mutation significantly reduced dsDNA binding by the β-domain (Fig. S5E) and abolished formation of the 90 kDa crosslinked species (Figs. 5D–E, S5D), supporting a role for the β-loop in bridging distal RAD51 protomers and dsDNA within RAD51–dsDNA filaments.

**Figure 5.**
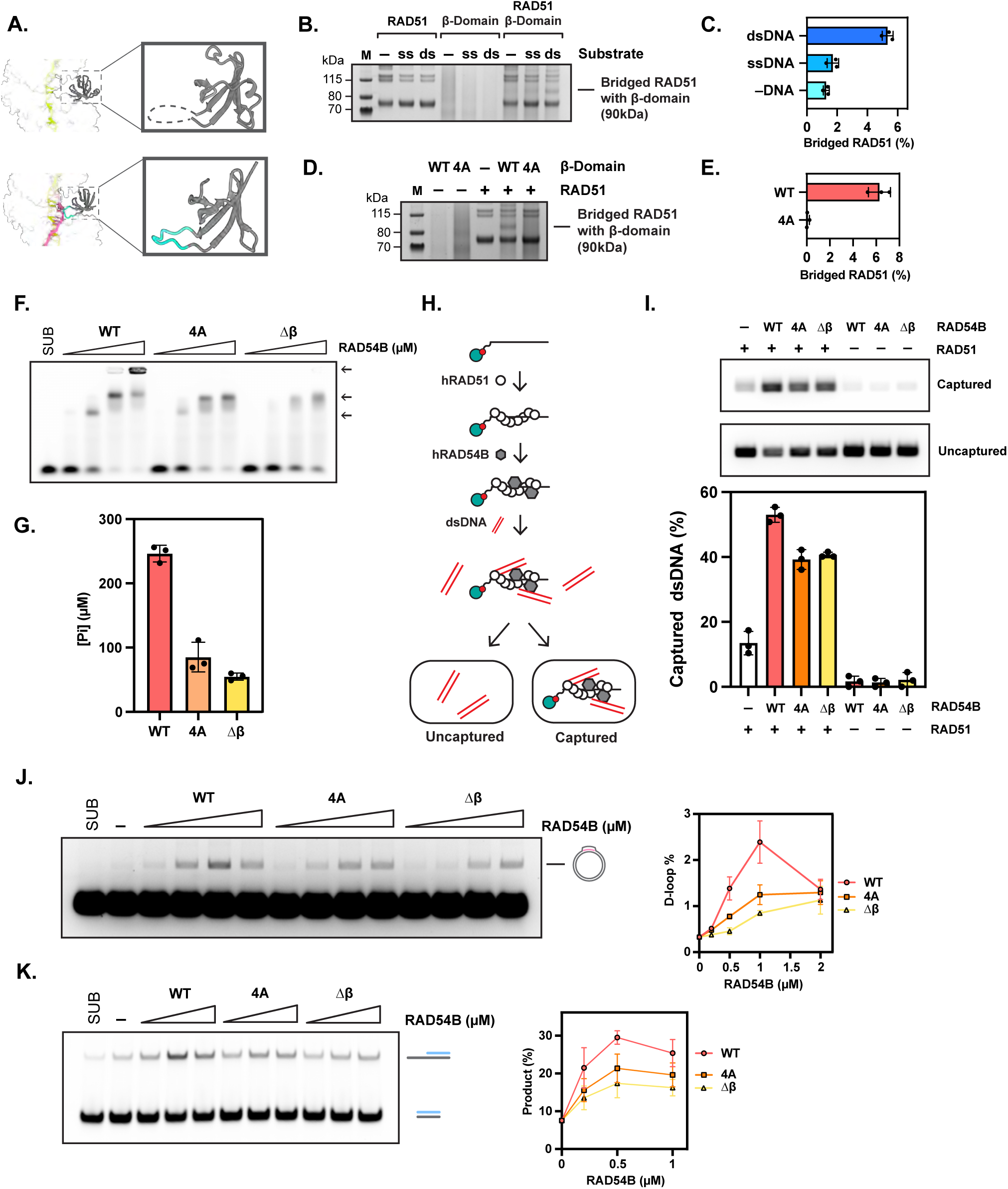
DNA-binding activity of the RAD54B β-domain is essential for maintaining intrinsic activity and bridging distal RAD51 protomers. **(A)** Comparison of the β-loop in the β-domain when bound to RAD51–ssDNA versus RAD51–dsDNA filaments. **(B)** Representing image gel of crosslinking experiment. RAD51, the β-domain, and mixtures of both in the absence or presence of ssDNA or dsDNA. Gel image cropped from Fig. S5C. **(C)** Quantification of **(B)**. **(D)** Crosslinking experiment of wild-type and 4A mutant β-domain with RAD51 in the presence of dsDNA. Gel image cropped from Fig. S7D. **(E)** Quantification of **(D)**. **(F)** Electrophoretic mobility shift assay (EMSA) showing binding of wild-type and mutant RAD54B (0.5, 1, 2, and 3 µM) to fluorophore-labelled 35-mer dsDNA. **(G)** ATPase assay comparing intrinsic ATPase activity of RAD54B (wild-type and mutants) in the presence of dsDNA. **(H)** Schematic of the dsDNA capture assay used to assess RAD51-mediated dsDNA capture. **(I)** Representative gel images showing stimulation of RAD51-dependent dsDNA capture by RAD54B (wild-type and mutants). **(J)** Representative gel images showing the stimulation of RAD51-driven D-loop formation by RAD54B (wild-type and mutants) (0.2, 0.5, 1, and 2 µM). **(K)** Representative gel images showing the stimulation of RAD51-driven strand exchange by RAD54B (wild-type and mutants) (0.2, 0.5, and 1 µM). For **(B-E, G, I-K)**, averages are shown with error bars depicting standard deviation (n = 3).

Given the strong contribution of the β-domain to D-loop formation and strand exchange (Figs. 4G–H), we next asked whether its dsDNA-binding activity plays a role in the context of full-length RAD54B. Introduction of the same 4A mutations into full-length RAD54B revealed that, although the isolated β-domain binds dsDNA weakly and this interaction is disrupted by the 4A mutation (Fig. S5E), neither the 4A mutation nor deletion of the β-domain significantly altered the overall dsDNA-binding affinity of full-length RAD54B (Fig. S5F). This is consistent with the presence of a conserved dsDNA translocase domain within the C-terminal motor region.

Despite these modest effects on bulk dsDNA binding, EMSA results revealed qualitative differences in complex formation. Full-length RAD54B formed three distinct dsDNA-bound species with a 35 bp substrate (Fig. 5F), suggestive of higher-order assemblies. Both the 4A mutant and the β-domain deletion mutant lacked the highest-molecular-weight species, indicating that the β-domain contributes to higher-order RAD54B–dsDNA assemblies. Consistent with this observation, both mutations caused a pronounced reduction in dsDNA-dependent ATPase activity (Fig. 5G), suggesting that the β-domain regulates RAD54B motor activity by influencing protein conformation and/or oligomeric state on dsDNA.

These findings imply that the β-domain, and specifically the positively charged loop, is required to maintain a functionally competent RAD54B–dsDNA complex. In light of this, we next tested whether the β-domain contributes to RAD54B-mediated bridging between RAD51 presynaptic filaments and donor dsDNA. In dsDNA capture assays, RAD51–ssDNA filaments immobilized on magnetic beads captured low levels of dsDNA on their own, whereas addition of RAD54B substantially enhanced dsDNA capture (Fig. 5I). This enhancement was reduced by approximately 25% in both the 4A mutant and the β-domain deletion mutant, indicating that the β-domain, and particularly the β-loop, contributes to RAD51-mediated dsDNA engagement.

Consistent with these defects, both the 4A mutant and β-domain deletion markedly impaired RAD54B stimulation of D-loop formation and strand exchange (Figs. 5J, K). Structural superposition of the RAD54B-RAD51-dsDNA complex with recently reported RAD51–D-loop structures (Greenhough et al. 2025; Luo et al. 2025; Joudeh et al. 2025) places the β-domain adjacent to the heteroduplex DNA within the D-loop, where it contacts the complementary strand in the homologous region during strand invasion (Figs. S5G–H). Together with the biochemical data, these observations support a role for the β-domain in promoting RAD51 filament conformational transitions from presynaptic filaments (ssDNA-bound) to synaptic filaments (D-loop intermediates) and ultimately to postsynaptic filaments (dsDNA-bound) (Fig. 5J; Fig. 6).

**Figure 6.**
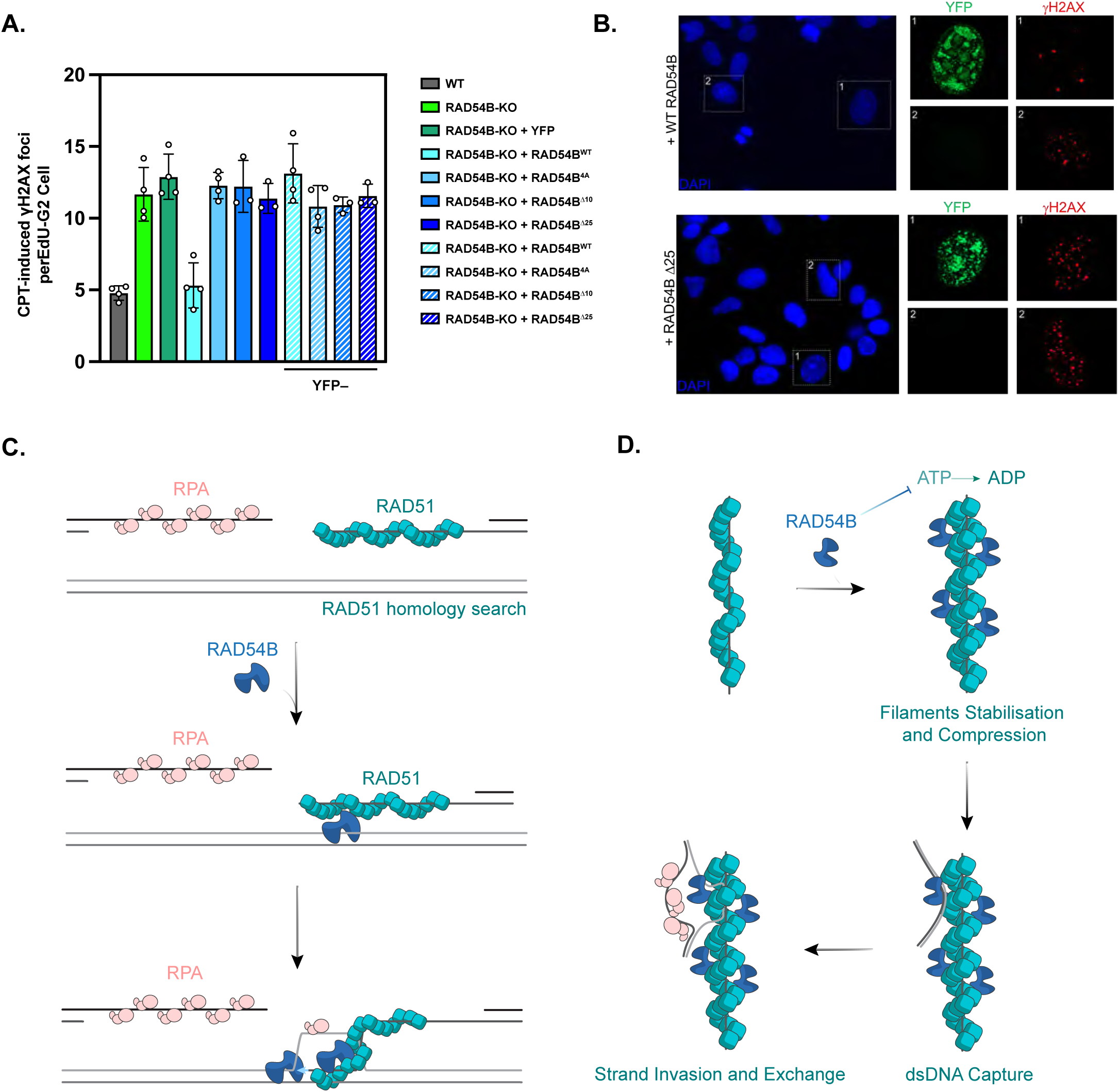
The RAD51 interaction sites and the β-domain of RAD54B are important for DSB repair by HR in U2OS cells. **(A)** γH2AX foci levels in U2OS WT and RAD54B-KO cells treated with 5 nM Camptothecin (CPT) for 24 h. RAD54B-KO cells were left untransfected or were transiently transfected with constructs for YFP-only, YFP-tagged RAD54B WT, RAD54B 4A, RAD54B Δ10 or RAD54B Δ25. 1 h before fixation, 10 µM EdU was added to monitor the cell cycle. γH2AX foci were quantified in EdU-negative G2-phase cells. In transfected cells, γH2AX foci were enumerated in YFP-positive (solid bars) and YFP-negative cells (striped bars). Spontaneous foci were subtracted (see Fig. S6). Bars represent the mean ± SEM (n=3-4), dots represent single experiments. **(B)** Representative IF images of γH2AX foci in cells treated with CPT. Overview images show the DAPI signal obtained from the scanning microscope. Cell nuclei with a G2 DNA content negative for EdU (not shown) and either positive (1) or negative (2) for YFP were selected. **(C)** Schematic model illustrating RAD54B binding to RAD51 filaments and donor dsDNA to stimulate subsequent homology pairing and recombination activities. **(D)** A structural basis for RAD54B’s functionalities in HR. Upon initial engagement with RAD51–ssDNA filaments, RAD54B stabilizes RAD51-ssDNA filaments, priming the filament for subsequent dsDNA capture. During homology search and pairing, the RAD54B motor domain functions as a DNA translocase to facilitate sequence sampling, while the β-domain can stabilize transient recombination intermediates via engaging the incoming dsDNA strand. Following homology recognition, RAD54B can stabilize and extend D-loop structure.

### The RAD51 interaction sites and the β-domain of RAD54B are important for DSB repair by HR in U2OS cells

To investigate the function of RAD54B in cells, we generated U2OS RAD54B-KO cells using CRISPR/Cas9 and validated them by sequencing. We treated exponentially growing cell cultures for 24 h with the topoisomerasere I inhibitor camptothecin (CPT). CPT induces single-stranded lesions which are converted during replication into DSBs. Following the progression of cells into G2 phase, such DSBs are repaired by HR (Ensminger and Lobrich 2020). Hence, we scored the level of γH2AX foci as a marker for DSBs in G2-phase cells as previously described (Lobrich et al. 2010). RAD54B-KO cells showed markedly elevated foci numbers compared with WT cells, indicating the importance of RAD54B for repairing CPT-induced DSBs (Figs. 6A, B and S5I). This finding is consistent with elevated γH2AX foci levels in RAD54B-deficient ovary cancer cells treated with a PARP inhibitor (Liu et al. 2022), which similar to CPT induces DSBs during S phase (Llorens-Agost et al. 2021). We then transiently transfected U2OS RAD54B-KO cells with YFP-tagged WT or mutated RAD54B constructs and assessed foci levels after CPT treatment in YFP-positive G2-phase cells (Fig. 6A). The expression of WT RAD54B reduced the foci levels of RAD54B-KO cells to those of WT cells. As a control, we analysed YFP-negative G2-phase cells in the same samples, which did not exhibit reduced foci numbers (Figs. 6B and S5I). In strong contrast to the expression of WT RAD54B constructs, the expression of RAD54B constructs carrying a RAD54B 4A, a RAD54B Δ10 or a RAD54B Δ25 mutation failed to reduce foci numbers in RAD54B-KO cells (Figs. 6B and S5I). This result shows that the two RAD51 interaction sites as well as the β-domain are important for RAD54B function in human cells.

## Discussion

### Unique contributions of the three sites and mechanisms of HR modulation

Our data presented here reveal that RAD54B acts as a multifunctional modulator that bridges RAD51 and dsDNA, stabilizing recombination intermediates and coordinating the transition from filament formation to strand invasion. The NTD mediates critical interactions with RAD51 and dsDNA, while the ATPase domain drives the DNA remodelling necessary for D-loop formation (Fig. 6C). The three sites identified in our structures have distinct contributions during HR. Binding experiments identify site 1 and site 2 to be responsible for recruiting RAD54B to RAD51-ssDNA filaments. Site 1 also inhibits RAD51 ATPase activity while sites 1 and β-domain (site 3) significantly contribute to filament stabilisation. ATPase inhibition stabilises RAD51-DNA filaments. However, filaments stabilisation does not necessarily result from ATPase inhibition as demonstrated by site 3, which stabilises the filaments by bridging two distal RAD51 protomers. Indeed, filaments can be stabilised by enhancing DNA binding and RAD51 protomer-protomer interactions. Inhibiting ATP hydrolysis stabilises DNA binding as ADP-bound form does not stably engage with DNA (Kuhlen et al. 2025a). Site 1 is located at the protomer-protomer interface and close to ATP binding site, therefore likely contributes to protomer-protomer interactions in addition to inhibit ATP hydrolysis. Site 3 contributes to RAD51-ssDNA filament stabilisation through bridging distal protomers and filament compression, which would also increasing protomer-protomer interface (Fig. 6D). In the dsDNA filament, site 3 also stabilises filaments via contacting dsDNA.

Since all sites contribute to strand exchange but only site 3 plays critical roles in D-loop formation, highlighting the differences in these two assays. Indeed, in oligonucleotides assays, strand invasion will promptly proceed to strand exchange while stable D-loop formation requires strand invasion, D-loop propagation and D-loop stabilisation due to tortional constraint imposed by plasmid DNA. In strand exchange assays, stable filaments are key to allow homology paring, and indeed strand invasion activities largely mirror that of filament stabilisation (Fig. 4F, 4J). In D-loop assays, deleting site 1 has no effects whereas deleting β-domain, although has no effects on RAD51 binding or RAD51 ATPase activity, is defective. These results support that RAD51 ATPase activities and filaments stabilisation only play limited roles in D-loop formation. In D-loop stabilisation requires ATPase activity of RAD54B which is regulated by β-domain (Figs. 4D, 4G, 5G).

Given that the β-domain interacts with dsDNA and influences dsDNA capture, it is plausible that this region contributes to donor dsDNA engagement during the homology search. Moreover, because the β-domain modulates RAD54B ATPase activity and higher-order assembly, it may help regulate the productive association of RAD54B with donor dsDNA, thereby enabling the motor activities required for D-loop formation. Consistent with structural comparisons to recently reported RAD51-D-loop (Greenhough et al. 2025; Luo et al. 2025; Joudeh et al. 2025) complexes that place the RAD54B β-domain adjacent to the heteroduplex DNA, where it contacts the complementary strand within the homologous region (Fig. S5G-H).

### RAD51 modulators use shared interaction sites on RAD51

RAD54B and RAD54 share conserved C-terminal ATPase domains but N-terminal domains differ. Our structure of RAD54B here identifies the β-domain that plays important roles in dsDNA binding, subsequently regulating the ATPase activity of RAD54B, its functional complex formation, D-loop and strand invasion activities. The β-domain is not conserved in RAD54, raising the question on how these activities in RAD54 are regulated. The N-terminal peptide is shown to be essential for RAD51 binding and inhibit RAD51 ATPase activity. Interestingly this region is highly conserved in RAD54 (Figs. S6A-E), and we suspected that it would bind to RAD51 in an identical fashion. Indeed the cryoEM structure of the RAD54 N-terminal peptide bound to RAD51 is identical to that of RAD54B and suggests this part of RAD54 might play similar roles (Figs. S6B-D). Additionally, the site 1 peptide that mediates RAD51 binding is highly conserved among higher eukaryotes but not in lower eukaryotes such as yeast, suggesting that this interaction has acquired increased functional importance during evolution (Figs. S6F-G).

The N-terminal 10-residue motif (site 1) of RAD54B and RAD54 represents a highly conserved RAD51-interacting element that is also shared by several other RAD51 regulators, including RAD51AP1, yeast Hed1, and yeast Rad54, suggesting that this region constitutes a major regulatory interface on RAD51 (Kuhlen et al. 2025a; Shin et al. 2025). Consistent with this idea, our recent work on RAD51AP1 demonstrated that this motif inserts into a deep cleft on the RAD51 surface, where it inhibits RAD51 ATPase activity and stabilizes RAD51 filaments. Yeast Hed1 and yeast Rad54 engage the same RAD51 surface, highlighting the evolutionary conservation of this interaction mode (Fig. S6H). In addition to this conserved N-terminal motif, a second RAD51-interacting element, the FxPP motif (site 2), is present in multiple RAD51 modulators, including the C-terminal TR2 motif of BRCA2, RAD54B, RAD51AP1, yeast Rad54, and FIGNL1 (Appleby et al. 2023; Longo et al. 2025; Kuhlen et al. 2025a; Carver et al. 2025; Shin et al. 2025). This motif primarily contributes to RAD51 binding (Fig. S6I). In the context of BRCA2 TR2, the FxPP motif has been shown to alter RAD51 dimer conformation and promote preferential binding to dsDNA (Longo et al. 2025). In contrast, our structures do not reveal substantial changes in RAD51 protomer interfaces upon RAD54B binding, suggesting that this motif can engage RAD51 in both filamentous and dimeric states without inducing major conformational rearrangements.

The shared sites between RAD51 modulators on RAD51 raise the question of if and how they might compete with each other for RAD51 binding, in particular for modulators that have overlapping activities such as stabilising RAD51 filaments or promoting D-loop formation. RAD54 and RAD54B are proposed to act synergistically on RAD51. Given that they share site 1, which is the primary RAD51 binding site, they are likely to bind to different RAD51 molecules within the filaments. It will be interesting to see how RAD54 and RAD54B, and indeed with other modulators, together interact and act on RAD51.

**Table 1.** Cryo-EM data collection and model statistics.

|  | RAD51 + RAD54B-N |  |  |  | RAD51 + RAD54-N |
| --- | --- | --- | --- | --- | --- |
| Nucleotide | ATP-Ca<sup>2+</sup> | ATP-Mg<sup>2+</sup> | ATP-Mg<sup>2+</sup> | ATP-Ca<sup>2+</sup> | ATP-Ca<sup>2+</sup> |
| DNA substrate | ssDNA | ssDNA | ssDNA | dsDNA | ssDNA |
| Microscope | Titan Krios |  |  |  | Glacios |
| Voltage (keV) | 300 |  |  |  | 200 |
| Detector | K3 (Gatan) |  |  |  | Falcon 4 |
| Magnification | 105,000 |  |  |  | 100,000 |
| Defocus range | -0.6 to -2 |  |  |  | -0.4 to -2 |
| Frames/movie | 60 |  |  |  |  |
| Pixel size | 0.84 |  |  |  | 1.14 |
| Electron dose | 60 |  |  |  |  |
| Exposure (s) | 3.2 |  |  |  |  |
| Movies | 6,086 | 25,595 | 25,595 | 14,860 | 8,876 |
| Picked particles | 5,171,621 | 1,690,118 | 1,690,118 | 2,317,245 | 1,754,084 |
| Final particles | 755,548 | 1,258,150 | 148,663 | 298,461 | 1,388,876 |
| Processing method | Single particle analysis | Single particle analysis | Single particle analysis | Single particle analysis | Single particle analysis |
| Resolution | 2.6 | 2.9 | 3 | 2.4 | 3.2 |
| Real-space refinement | Non-uniform Refinement | Homogenous Refinement | Non-uniform Refinement | Homogenous Refinement | Helical Refinement |
| **Composition** |  |  |  |  |  |
| Non-H atoms | 15809 | 15853 | 16210 | 16602 | 15361 |
| Protein residues | 1986 | 1986 | 2031 | 2047 | 1926 |
| Nucleotide residues | 18 | 20 | 20 | 35 | 18 |
| Ligands | 12 | 17 | 17 | 18 | 18 |
| Crosscorrelation (CC, mask) | 0.9 | 0.87 | 0.85 | 0.91 | 0.89 |
| **Bonds (rmsd)** |  |  |  |  |  |
| Lengths (Å) | 0.002 | 0.003 | 0.004 | 0.003 | 0.003 |
| Angles ( ° ) | 0.464 | 0.532 | 0.992 | 0.732 | 0.525 |
| MolProbity score | 1.07 | 1.36 | 1.56 | 1.25 | 1.49 |
| Clash score | 2.64 | 6.13 | 8.13 | 4.8 | 9.28 |
| Rotamer outliers (%) | 1.07% | 1.08% | 1.42% | 0.61% | 0.71% |
| **Ramachandran plot** |  |  |  |  |  |
| Outliers (%) | 0.00% | 0.00% | 0.00% | 0.00% | 0.00% |
| Allowed (%) | 0.52% | 1.34% | 1.87% | 1.56% | 1.49% |
| Favoured (%) | 99.48% | 98.66% | 98.13% | 98.44% | 98.51% |
| **B factors** |  |  |  |  |  |
| Protein | 29.38 | 52.57 | 70.31 | 36.65 | 41.35 |
| Nucleotide | 30.16 | 52.59 | 52.28 | 29.09 | 42.62 |
| Ligand | 31.86 | 60.23 | 67.84 | 38.24 | 44.25 |

## Resource Availability

## Acknowledgement

We thank members of the Zhang lab for their helpful insights and discussions. Initial screening of electron microscopy grids was carried out at Imperial College London Centre for Structural Biology. We acknowledge Diamond Light Source for access and support of the cryoEM facilities at the UK national electron bioimaging centre, proposal B136390, funded by the Wellcome Trust and the United Kingdom Research and Innovation Medical Research Council, and London Consortium for high resolution cryoEM (LonCEM), funded by the Wellcome. This work is funded by the Wellcome Trust to X.Z. (210658/Z/18/Z, 227769/Z/23/Z); the National Institutes of Health to WDH (R01GM58015) and the Deutsche Forschungsgemeinschaft to ML (DFG, German Research Foundation – Project-ID 393547839 – SFB 1361).

## Author Contributions

P.L. and X.Z. designed the study. P.L., S.T., M.B. and E.M. purified proteins. P.L. and S.T. carried out biochemical and biophysical analysis. N.M. conducted the ATPase experiment of RAD54B on RAD51-bound DNA. P.L. performed EM studies and data analysis. J.E. conducted the cell biology experiments. B.A. and L.K. contributed on methodology. X.Z. M.L and W.D.H. supervised the experiments and data analysis. P.L., S.T., B.A., J.E., M.L., W.D.H. and X.Z. prepared the manuscript with contributions from all of the authors.

## Declaration of interests

Authors declare that they have no competing interest.

## Declaration of generative AI and AI-assisted technologies

AlphaFold3 server (Abramson et al., 2024) was used to generate initial structural model of RAD54B which was modified and refined according to experimental data. EMReady (He et al., 2023) was used for density modification of experimental EM maps after data processing.

## Supplemental Information

Figures S1-6

## STAR Methods

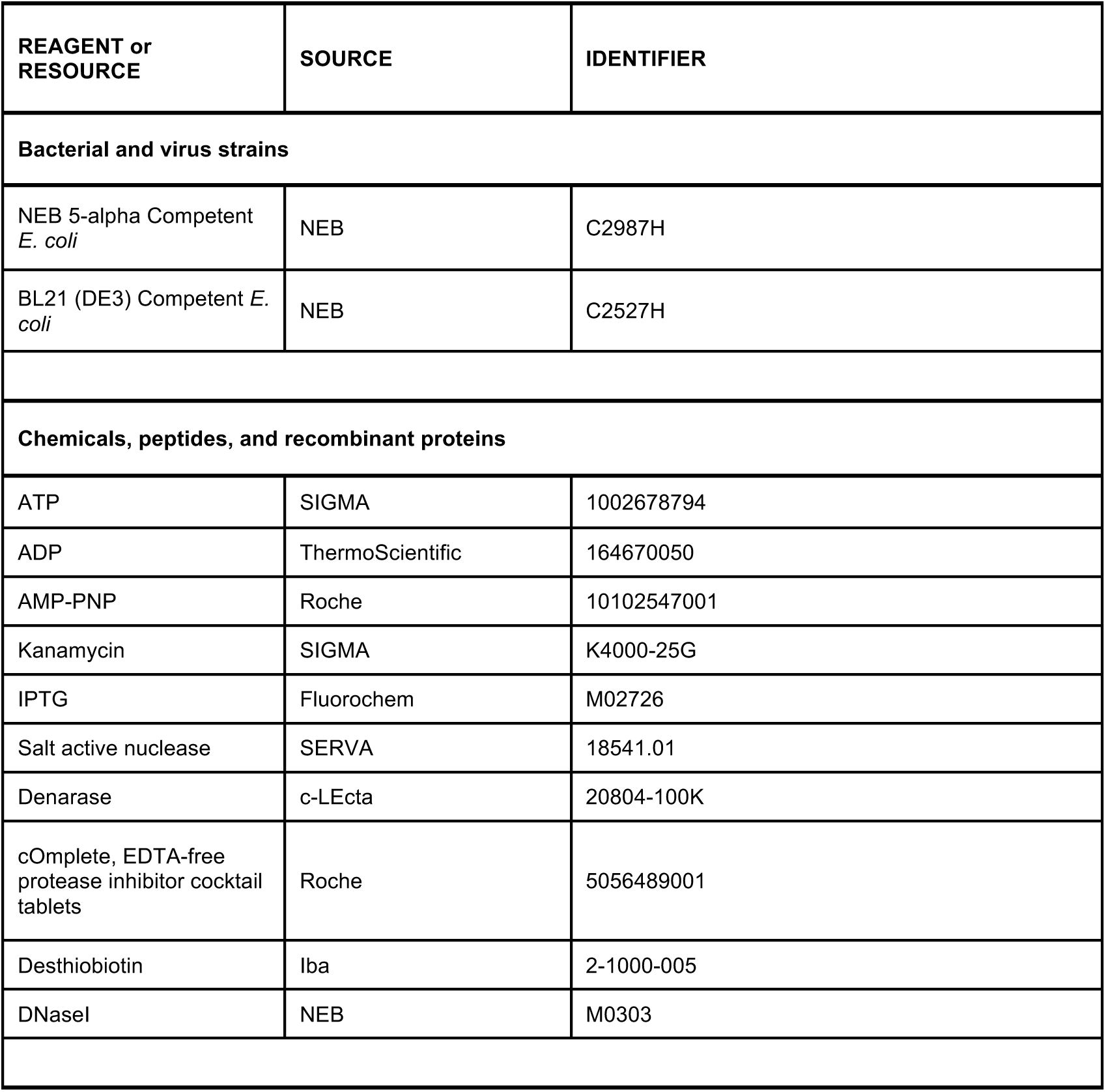
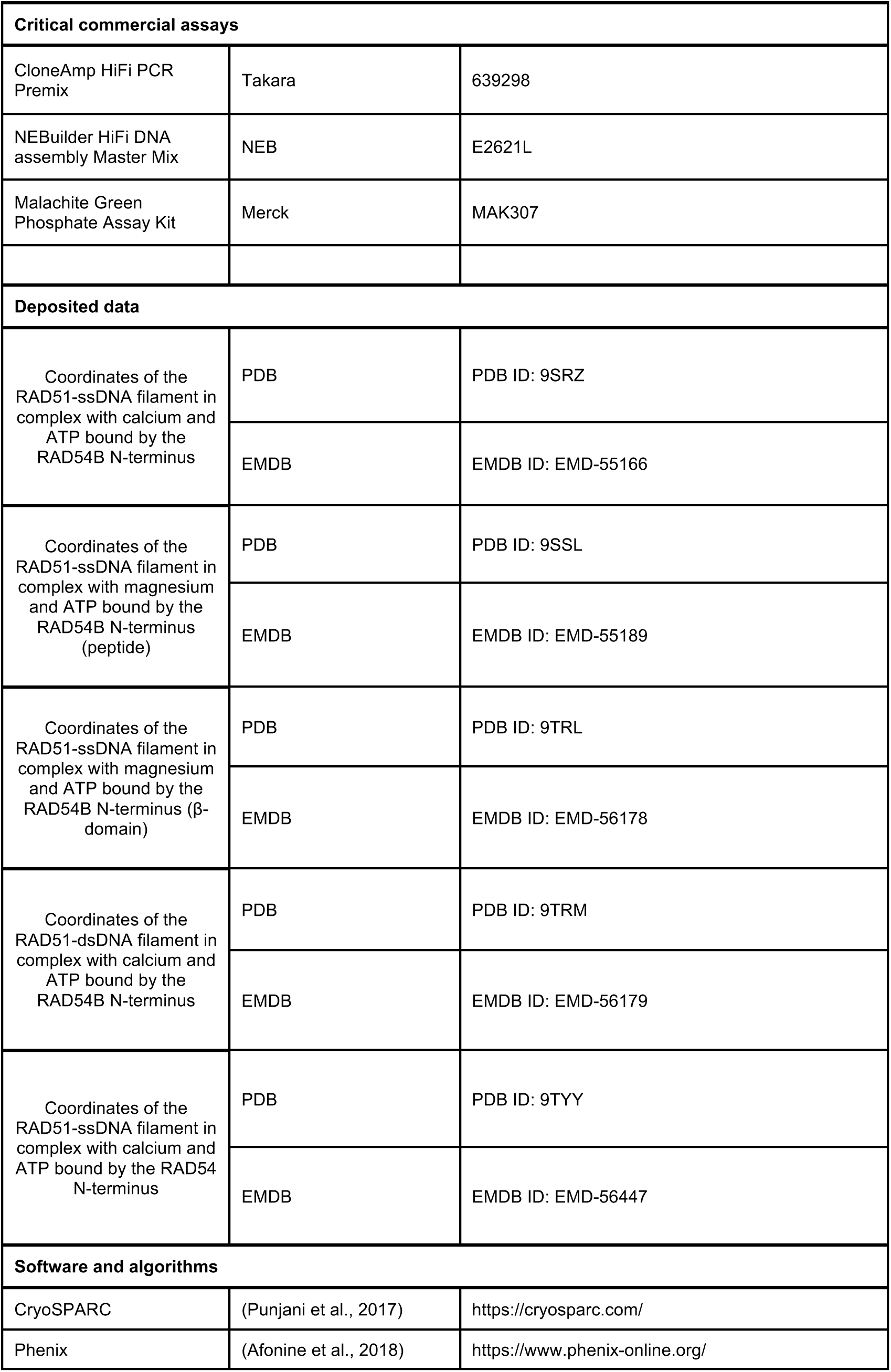
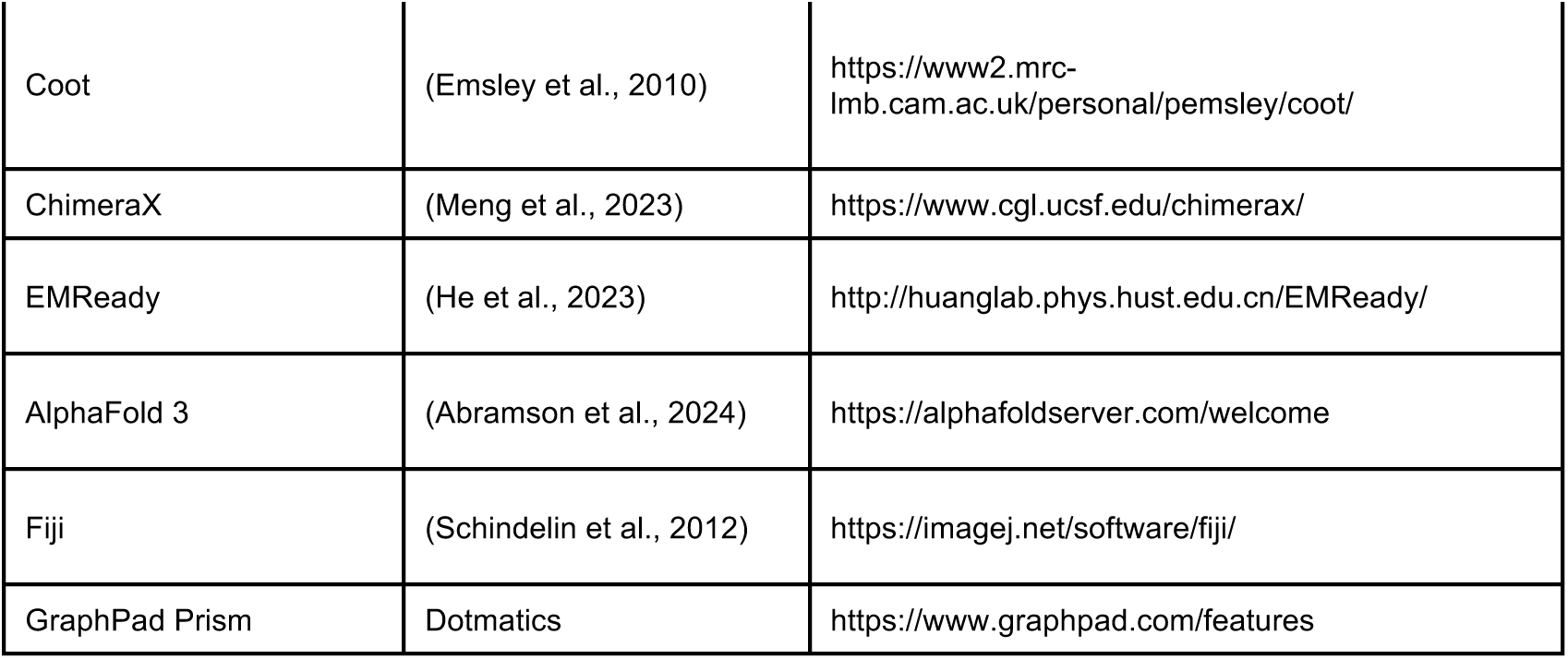

**Oligonucleotides**

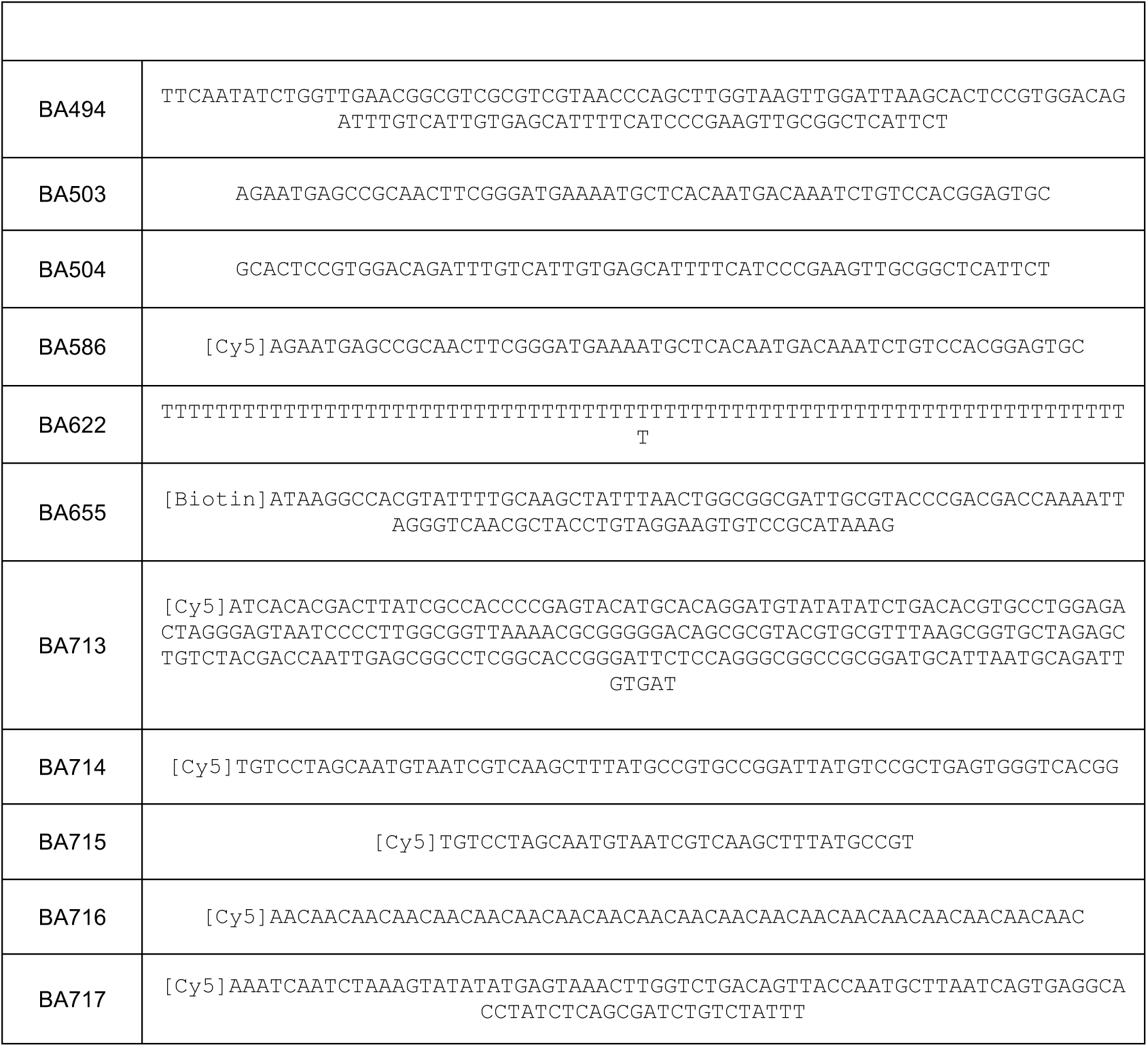

### Sequence Alignment

A multiple sequence alignment of RAD54B sequences of representative species was generated using Clustal Omega (Sievers and Higgins 2018).

### Cloning

The Human RAD54B with a C-terminal Twin-Strep (TS) or 10xHis-Twin-Strep tag sequence was synthesized from IDT, amplified by PCR, and inserted into a pFBDM vector. Substitution and truncations were generated by PCR, and the fragments were combined using the NEBuilder HiFi assembly kit (NEB).

### Protein Purification

Human RAD51 was cultured and purified as previous descried (Kuhlen et al. 2025a). RAD51^C319S^ was cultured and purified in the same way as wild-type protein. To label the RAD51^C319S^, 15 µL of AuroraFluor 488 (Biosynth Laboratories Ltd, 25 mM in DMSO) was incubated with 1.3 mL RAD51^C319S^ at 1.12 mg/mL at 4°C overnight. Then further purification was carried on HiTrap Heaprin HP column in 20mM Tris [pH7.5], 150-2000 mM NaCl, 0.5 mM TCEP.

The Full-length RAD54B, mutants, and RAD54-N/C proteins were expressed in baculovirus-infected Hi5 cells. Hi5 cells (density 1 x 10^6^) were infected with the recombinant baculovirus at multiplicity of infection of 2 and after 48 h of incubation were collected by centrifugation (1000 x g). Cells pellets were resuspended in lysis buffer (20 mM HEPES [7.5], 300 mM KCl, 0.5 mM TCEP, 10% glycerol) supplemented with cOmplete protease inhibitors (Roche^TM^). Cells were lysed by sonication and clarified by centrifugation before applying to a 5 mL StrepTrap HP column (Cytiva^TM^). The column was washed by 50 mL lysis buffer and eluted in 50 mL elution buffer (20 mM HEPES [7.5], 200 mM KCl, 0.5 mM TCEP, 10% glycerol, 10 mM desthiobiotin). The protein was further purified using a 5 mL Heparin HP column (Cytiva^TM^) run in RA buffer (20 mM Tris [pH7.5], 0-1 M KOAC, 0.5 mM TCEP, 10% glycerol). The peak fractions were pooled and concentrated to 0.5 mL before loading to a Superdex 200 increase 10/300 column (Cytiva^TM^) equilibrated with RA150 buffer (20 mM Tris [7.5], 150 mM KOAC, 0.5 mM TCEP, 10% glycerol). Finally, the proteins were concentrated and frozen in aliquots in liquid nitrogen.

RAD54B beta-domain (wild-type and the mutant) was expressed from Hi5 cells with an N-terminal 6xHis tag and a C-terminal Twin-Strep tag. RAD54B beta-domain (WT) purified in a similar scheme as RAD54B Full-length purification except the gel filtration column is Superdex 75 increase 10/300 (Cytiva^TM^). RAD54B beta-domain (4A mutant) is initially bound and eluted from a 5mL StrepTrap HP column. Then loaded on a 5mL HiTrap Q column (Cytiva^TM^). The flow-through from Q column is collected before applying to a 5mL HisTrap HP column (Cytiva^TM^) equilibrated with H150 buffer (25 mM HEPES [7.5], 150 mM NaCl, 5% glycerol, 0.5 mM TCEP), then eluted in 40 mL H150 supplemented with 300mM Imidazole. The proteins were concentrated, and gel filtrated with Superdex 75 increase 10/300 column.

### Pull-down assays

Twin-Strep- or 10×His-Twin-Strep-tagged RAD54B (1 µM) in 30 µL of pull-down buffer (30 mM Tris-OAc [pH 7.5], 100 mM KCl, 2 mM ATP, 0.1% Tween-20, 1 mM TCEP, and 5% glycerol) was immobilized on Strep-Tactin Sepharose High Performance resin (Cytiva™) by incubation at 25 °C for 30 min with mixing. RAD51 or RAD51 (C319S)-AF488 (1 µM) was then added and incubated for an additional 30 min with mixing. The supernatant containing unbound RAD51 was collected, and bound proteins were eluted by incubation with SDS-PAGE loading buffer (65 °C, 1300 rpm, 10 min). Protein content in both fractions was analysed by SDS-PAGE, and gels were imaged using a Bio-Rad ChemicDoc MP Imaging System™. Band intensities were quantified using FIJI. Images were background-subtracted using the rolling ball method (50 pixels), and the intensities of RAD51 and RAD54B bands in the supernatant and eluate were measured. The percentage of protein recovered in the eluate was then calculated.

### ssDNA Nuclease Protection Assay

A Cy5-labeled oligonucleotide (60-mer, BA586; 10 µMnt) in ssDNA nuclease protection buffer (25 mM Tris-OAc [pH 7.5], 100 mM KCl, 2 mM ATP, 5 mM MgCl₂, 0.5 mM TCEP, 2.5% glycerol) was incubated with RAD51 (1 µM) at 37 °C for 5 min prior to addition of the indicated RAD54B. After a further 5 min incubation at 37 °C, DNase I (4 units; NEB™ M0303) was added and reactions were incubated for 10 min. Reactions were stopped by addition of deproteinization buffer (final concentrations: 12 mM Tris-Cl [pH 7.5], 7.5 mM EDTA, 0.45% SDS, 0.75 mg/mL proteinase K). DNA products were resolved on 10% PAGE gels and imaged using a Bio-Rad ChemiDoc MP Imaging System™. Quantification was performed in FIJI (Schindelin et al., 2012). Images were background-subtracted using the rolling ball method (50 pixels), and the total lane signal was measured. The fraction of signal corresponding to intact DNA was expressed as a percentage of the total.

### dsDNA Nuclease Protection Assay

A Cy5-labeled oligonucleotide (213-mer, BA713; 5 µMbp) in dsDNA nuclease protection buffer (25 mM Tris-OAc [pH 7.5], 50 mM KCl, 2 mM ATP, 5 mM MgCl₂, 0.5 mM TCEP, 2.5% glycerol) was incubated with RAD51 (0.5 µM) at 37 °C for 5 min prior to addition of the indicated RAD54B. After a further 5 min incubation at 37 °C, DNase I (2 units; NEB™ M0303) was added and reactions were incubated for 10 min. Reactions were stopped by addition of deproteinization buffer (final concentrations: 12 mM Tris-Cl [pH 7.5], 7.5 mM EDTA, 0.45% SDS, 0.75 mg/mL proteinase K). DNA products were resolved on 2% agarose gels and imaged using a Bio-Rad ChemiDoc MP Imaging System™. Quantification was performed in FIJI. Images were background-subtracted using the rolling ball method (50 pixels), and the total lane signal was measured. The fraction of signal corresponding to intact DNA was expressed as a percentage of the total.

### D-loop Assay

RAD51 (1 µM) was incubated with a 90-mer ssDNA oligonucleotide (BA717; 6 µMnt) in D-loop buffer (25 mM Tris-OAc [pH 7.5], 50 mM KCl, 2 mM ATP, 2 mM MgCl₂, 0.5 mM CaCl₂, 0.5 mM TCEP, 0.1 mg/mL BSA, 3 mM phosphocreatine, 20 u/mL CPK) at 37 °C for 5 min. The reaction was initiated by adding the indicated RAD54B and pBlueScript (pBS-SK-II; 120 µMnt). Reactions were incubated at 37 °C for 10 min, deproteinized, resolved by PAGE, and imaged as described above. Quantification was performed using FIJI (Schindelin et al. 2012). Images were background-subtracted using the rolling ball method (50 pixels), and the total lane signal was measured. The fraction of signal corresponding to product was expressed as a percentage of the total. For each experiment, the value for the substrate alone was subtracted from all samples.

#### Strand Exchange Assay

RAD51 (1 µM) was incubated with a 116-mer ssDNA oligonucleotide (BA494; 3 µMnt) in strand exchange buffer (25 mM Tris-OAc [pH 7.5], 50 mM KCl, 2 mM ATP, 1 mM MgCl₂, 2 mM CaCl₂, 0.5 mM TCEP, 0.1 mg/mL BSA) at 37 °C for 7 min. RPA (0.2 µM) was then added, and after an additional 7 min incubation at 37 °C, the reaction was initiated by adding the indicated RAD54B and 60-mer dsDNA (BA586-BA504; 1 µM bp). Reactions were incubated at 37 °C for 10 min, deproteinized, resolved by PAGE, and imaged as described above. Quantification was performed using FIJI (Schindelin et al. 2012). Images were background-subtracted using the rolling ball method (50 pixels), and the total lane signal was measured. The fraction of signal corresponding to product was expressed as a percentage of the total. For each experiment, the value for the substrate alone was subtracted from all samples.

### ATPase Assay

All of ATPase assays were conducted using a malachite green reagent-based spectrophotometric method (Ehmsen et al. 2019) This is a sensitive method for the detection of released phosphate, in which a green complex formed between Malachite Green, molybdate, and free orthophosphate is measured on a spectrophotometer at 620nm.

RAD51 (3 µM) and poly-dT ssDNA (BA622, 72-mer; 9 µMnt) were incubated in ATPase buffer (35 mM Tris-OAc [pH 7.5], 50 mM KCl, 0.5 mM ATP, 2 mM MgCl_2_, 1 mM TCEP, 5% glycerol) supplemented with RAD54B (3 µM) at 37 °C for 60 min. Reactions were stopped by addition of 50 mM EDTA and diluted two-fold to reduce phosphate concentration. Absorbance was measured at 620 nm using a CLARIOstar plate reader, and phosphate concentration was determined with a commercial colorimetric kit (MAK307, Merck) according to the manufacturer’s instructions.

RAD54B intrinsic ATPase activity was measured by incubating 150 nM RAD54B and 30 µMnt pBS-SK-II plasmids in ATPase buffer but with 0.25 mM ATP (37°C, 10 min). Reactions were stopped by addition of 50 mM EDTA and diluted 20-fold to reduce phosphate concentration. Absorbance was measured at 620 nm using a CLARIOstar plate reader, and phosphate concentration was determined with a commercial colorimetric kit (MAK307, Merck) according to the manufacturer’s instructions.

To measure the effects of RAD51 (Sigurdsson et al. 2001) on RAD54B ATPase activity, ATP hydrolysis assays were performed using a two-solution reaction setup. Solution I contained 25 mM HEPES (pH 7.5), 7 mM MgOAc₂, 1 mM DTT, 30 µg/mL BSA, 0.1% CHAPS, 2 mM ATP, with or without ΦX174 RFI dsDNA (12 µM bp; New England Biolabs) and with or without RAD51 (300 nM). Solution II contained the same buffer components and ATP concentration, with or without RAD54B (20 nM; full-length or mutant variants). Ten microliters of Solution I containing dsDNA with or without RAD51 (1 RAD51 per 40 bp) were incubated at 37 °C for 10 min, after which 10 µL of Solution II containing RAD54B full-length or mutant proteins (Δ25 or Δβ-domain; 1 RAD54B per 600 bp or 1 RAD54B per 15 RAD51 molecules) was added and incubated for an additional 10 min at 37 °C. Reactions were developed by adding 6 µL of malachite green reagent (1.6% ammonium molybdate, 0.16% malachite green, 4.26 M HCl, 1.07% polyvinyl alcohol), followed immediately by quenching with 6 µL of 6% sodium citrate, bringing the final reaction volume to 32 µL. Absorbance was measured at 620 nm using a SpectraMax iD5 plate reader (Molecular Devices). Reactions lacking protein were used as blank and negative controls. ATP hydrolysis rates were calculated using the formula “Rate of ATP hydrolysis = {Fraction of normalized values (NV) x [ATP] (nM)}/ time of reaction (min-1)/ concentration of protein(nM)}”.

### Electrophoretic Mobility Shift Assay

RAD51 (5 µM) and ssDNA (BA714, 65-mer; 15 µMnt) or dsDNA (BA586-504, 60-mer; 12 µMbp) were incubated in filaments EMSA buffer (25 mM HEPES-KOH [pH 7.5], 1 mM TCEP, 100 mM KCl, 5 mM MgCl₂, 2 mM ATP) at 25 °C for 10 min, followed by addition of the indicated RAD54B and incubation for 5 min. Reactions were crosslinked with 0.1% glutaraldehyde for 5 min and quenched with 1 M Tris-Cl [pH 7.5]. Products were resolved on 0.8% agarose gels.

The indicated RAD54B and dsDNA substrates (BA715, 35-mer; 17.5 µMbp) were incubated in binding buffer (25 mM Tris-OAc [pH 7.5], 1 mM TCEP, 100 mM KOAc, 5 mM MgOAc₂, 0.1 µg/µL BSA) at 25 °C for 10 min. Products were resolved on 0.8% agarose gels.

### RAD51-Bridge Crosslink Assay

RAD51 (6 µM) was incubated with or without DNA substrates (BA716, 60-mer, 18 µMnt; BA503-504, 60-mer, 18 µMbp) in crosslinking buffer (25 mM HEPES-KOH [pH 7.5], 1 mM TCEP, 2 mM ATP, 5 mM CaCl₂) at 37 °C for 5 min. RAD54B β-domain (15 µM) was then added, and reactions were incubated for an additional 15 min. Crosslinking was initiated by adding 5 mM bis(sulfosuccinimidyl) suberate, followed by 5 min incubation at room temperature. Products were resolved by SDS–PAGE.

### dsDNA Capture Assay

Biotin-labeled ssDNA (BA655, 100-mer; 5 µMnt) in capture buffer (25 mM Tris-OAc [pH 7.5], 1 mM DTT, 150 mM KOAc, 0.05% Tween-20, 2 mM ATP, 5 mM MgOAc₂, 5% glycerol) was immobilized on magnetic beads (Dynabeads™ M-280 Streptavidin) prior to addition of RAD51 (1.67 µM) and incubation at 25 °C for 10 min with shaking. RAD54B (1 µM) was then added and incubated for an additional 5 min. The supernatant was removed using a magnetic stand, and the bead-bound complexes were resuspended in 10 µL capture buffer containing fluorophore-labeled dsDNA (BA586-504, 60-mer; 5 µMbp) and incubated with mixing at 25 °C for 30 min. The supernatant containing uncaptured dsDNA was collected, and bead-bound dsDNA was eluted by deproteinization (37 °C, 15 min, 1200 rpm). DNA from both fractions was analyzed by agarose gel electrophoresis.

### MST

RAD54B-dsDNA interaction studies (full-length or β-domain) were performed using 5′ Cy5-labeled dsDNA substrates (35-mer, BA715). Fluorescently labeled DNA in binding buffer (25 mM Tris-OAc [pH 7.5], 1 mM TCEP, 50 mM KOAc, 0.05% Tween-20) was diluted to 10 nM and mixed with an equal volume of a serial dilution of RAD54B. Samples were incubated at room temperature for 15 min before loading into MST capillaries. MST experiments were carried out on a Monolith X instrument (NanoTemper Technologies) using premium-treated capillaries at 25 °C. Binding data were analyzed using Monolith software and replotted in Prism.

### Negative Stain

Samples were taken from filaments EMSA (with 1µM RAD54B) followed by 2 times dilution before applied to glow discharged 300 mesh carbon coated copper grids (Agar Scientific), blotted and stained with 2% uranyl acetate. For each sample, ∼200 micrographs were collected on a T12 microscope (FEI) at a magnification of 30,000 using a Rio CMOS camera (Gatan^TM^). Then micrographs were processed in cryoSPARC for 2D classification.

### Cryo-EM Sample Preparation

RAD51 (2 µM) was incubated with ssDNA (BA716; 6 µMnt) or dsDNA (BA503-504; 6 µMbp) in filament EMSA buffer (MgCl₂ replaced with CaCl₂ where indicated) at 25 °C for 10 min to form filaments. Filament samples (4 µL) were applied to plasma-cleaned 300-mesh lacey carbon grids (LC300-Au-UL, EM Resolutions). Next, 3 µL of filament sample was replaced with 3 µL of RAD54B-N or RAD54-N (10 µM). Grids were then blotted and plunged into liquid ethane using a Vitrobot Mark IV (FEI) with a blotting force of -2, a waiting time of 90 s, and a blotting time of 2 s.

### Cryo-EM Data Processing

For all datasets, movies were corrected with MotionCor2 as implemented in RELION5 (Zivanov, Nakane, and Scheres 2019). All subsequent processing was performed in cryoSPARC versions 4.0–4.6 (Punjani et al. 2017).

### RAD54B-N bound to RAD51 ssDNA filaments (Ca^2+^)

A total of 6,082 micrographs were imported into cryoSPARC, and CTF parameters were estimated using PatchCTF. Approximately 5 million particles were initially picked using a combination of template-free filament tracing and template picker, with 2D class averages of RAD51 filaments from a previous screening dataset as templates. Particles were extracted with a box size of 320 × 320 pixels and fourfold binned for initial processing. Following 2D classification, ∼1 million particles were re-extracted with the same box size and twofold binning. These particles were subjected to 3D reconstruction with template-free helical refinement, using an estimated helical twist of 56° and rise of 16 Å. Refinement optimized these parameters to 55.9° and 16.2 Å. Duplicate particles were removed, and the remaining set was re-extracted at the same box size without binning. In total, 755,548 particles and the 3D volume from helical refinement were used in non-uniform refinement, yielding a 2.6 Å reconstruction. Map interpretability was further improved with EMReady (He, Li, and Huang 2023).

### RAD54B-N bound to RAD51 ssDNA filaments (Mg^2+^)

A total of 29,547 micrographs were imported into cryoSPARC, and CTF parameters were estimated with PatchCTF. Approximately 1.6 million particles were picked using a combination of template-free filament tracing and template picker, with 2D averages of RAD51 filaments as templates.

For the β-domain–bound class, particles were extracted with a box size of 360 × 360 pixels and fourfold binned. 3D reconstruction was performed with template-free helical refinement (initial twist 56°, rise 16 Å), which refined to 55.5° and 16.3 Å. The refined model was then used for homogeneous refinement, followed by three rounds of 3D classification with a loose mask around the filament groove. A subset of 148,663 particles was re-extracted without binning and refined with non-uniform refinement, yielding a 3.0 Å reconstruction.

For the peptide–bound class, particles were extracted with a 360 × 360 pixel box size and twofold binned. These were processed by homogeneous refinement, duplicate particles removed, and the remainder re-extracted without binning. A total of ∼1.2 million particles and the 3D volume from helical refinement were used in non-uniform refinement, yielding a 2.9 Å reconstruction. Map interpretability in both cases was further improved using EMReady (He, Li, and Huang 2023).

### RAD54B-N bound to RAD51 dsDNA filaments (Ca^2+^)

A total of 14,860 micrographs were imported into cryoSPARC, and CTF parameters were estimated with PatchCTF. Approximately 2.3 million particles were picked and extracted with a 360 × 360 pixel box size and fourfold binned for initial 2D classification. After classification, particles were re-extracted at twofold binning, split into two subsets, and subjected to template-free helical refinement (initial twist 56°, rise 16 Å). The refined model was used for homogeneous refinement, followed by two rounds of 3D classification with a loose mask around the filament groove. Selected particles were re-extracted without binning, followed by an additional 3D classification and homogeneous refinement, resulting in a 2.4 Å reconstruction. Map interpretability was improved with EMReady (He, Li, and Huang 2023).

### RAD54-N bound to RAD51 ssDNA filaments (Ca^2+^)

A total of 8,876 micrographs were imported into cryoSPARC, and CTF parameters were estimated with PatchCTF. Approximately 6 million particles were picked, extracted with a 440 × 110 pixel box size, and fourfold binned. Following 2D classification, ∼1.7 million particles were re-extracted without binning. These particles were subjected to template-free helical refinement (initial twist 56°, rise 16 Å), which refined to 56° and 15.6 Å. Duplicate particles were removed, and 1,388,876 particles and the refined 3D volume were subjected to another round of helical refinement, yielding a 3.2 Å reconstruction. Map interpretability was improved with EMReady (He, Li, and Huang 2023).

### Model Building

Structural models comprising six RAD51 protomers bound to ssDNA or dsDNA, ATP, Ca²⁺ or Mg²⁺, and RAD54B-N or RAD54-N (as appropriate) were generated using AlphaFold (Abramson et al. 2024a). These models were rigid-body docked into the cryo-EM maps, manually adjusted in Coot (Emsley et al. 2010), and subsequently refined in Phenix (Afonine et al. 2018).

### Cell Biology experiments

#### Cell lines and cell culture

U2OS WT and RAD54B-KO cells were cultured at 37°C with 5% CO_2_ in Dulbecco’s Modified Eagle’s Medium (DMEM, Sigma) supplemented with 10% fetal calf serum (FCS), 1% Penicillin/Streptomycin, and 1% non-essential amino acids.

#### U2OS RAD54B-KO generation using CRISPR/Cas9

U2OS cells were transfected with pSP-Cas9(BB)-2A-Puro (PX459) V2.0 carrying the sgRNA 5’-CATCTAATATGTATAGGAGC-3’. 24 h later, medium was changed to selection medium containing 1 µg/ml Puromycin. After nine days, single cells were seeded to obtain colonies. Genomic DNA was isolated using DNeasy Blood and Tissue Kit (Qiagen). The target region was amplified using Q5 Polymerase and the following primers: fwd 5’-ACTTACTTGCCTCACTGGACTC-3’ and rev 5’-TAAGGTGCCAAAGGGGATT-GG-3’. The knockout was verified by sequencing. For allele 1, sequencing revealed a 2 bp deletion and therefore a frameshift starting at aa581, resulting in a premature stop codon at aa588. Allele 2 showed an insertion of 171 bp at aa582, resulting in a premature stop codon after 15 of the inserted basepairs. pSpCas9(BB)-2A-Puro (PX459) V2.0 was a gift from Feng Zhang (Addgene plasmid # 62988; http://n2t.net/addgene:62988; RRID: Addgene_62988).

#### Site-directed mutagenesis

A vector encoding the human RAD54B sequence C-terminally tagged with YFP was purchased from Vectorbuilder and subjected to site-directed mutagenic PCR (primer sequences to introduce the different mutations are listed in the table). Three different mutations were induced: a RAD54B 4A mutant carries four subsequent mutations: K135A, K136A, K137A and H138A. A RAD54B Δ10 mutant contains a deletion of aa2-10, and a RAD54B Δ25 mutant contains a deletion of aa2-25. The plasmid carrying the wt RAD54B sequence was amplified by PCR using either the fwd or rev primer of the desired mutation. The reactions were then pooled, realigned and digested using DpnI restriction enzyme. After transformation into VB UltraStable E. Coli (Vectorbuilder), clones were validated by colony PCR and whole-plasmid sequencing.

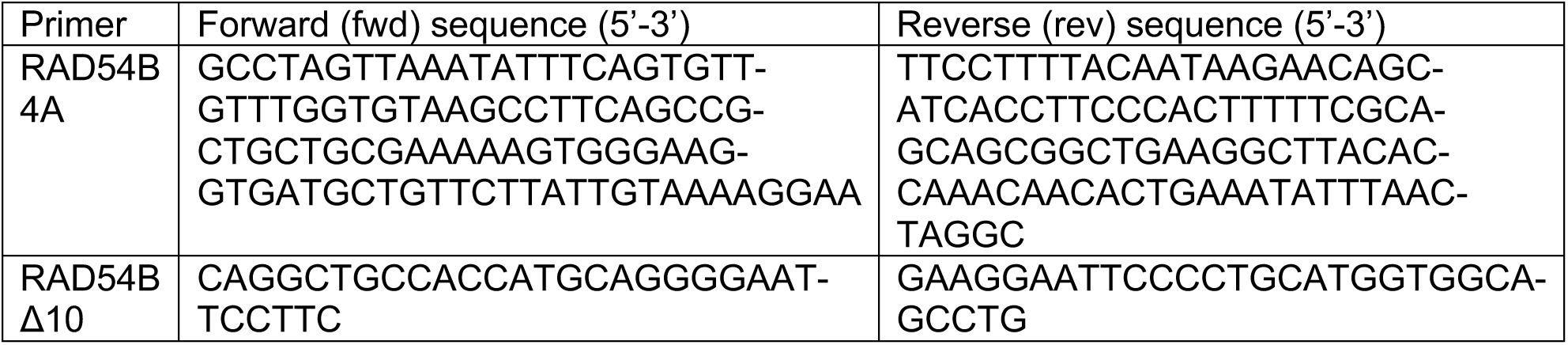
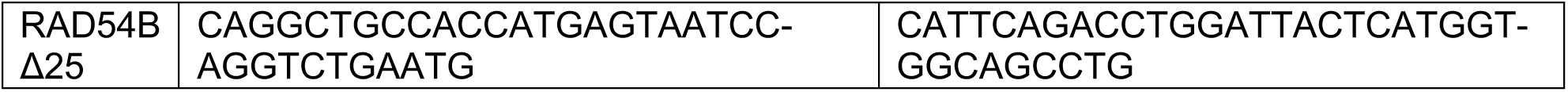

#### Plasmid transfection

U2OS RAD54B-KO cells were transfected with different plasmids (pRP-EYFP, pRP-RAD54B-EYFP, pRP-RAD54B-4A-EYFP, pRP-RAD54B-Δ10-EYFP and pRP-RAD54B-Δ25-EYFP) using Lipofectamine 2000 (Thermo Fisher Scientific) according to manufacturer’s instructions. Experiments were performed 48 h after transfection.

#### Immunofluorescence

Cells were cultured on glass coverslips and fixed with 2.5% formaldehyde in PBS for 15 min at room temperature (RT), washed with PBS-Tween (PBS-T) and permeabilized with 0.2% Triton-X 100 in PBS-T/1% FCS for 5 min at 4°C. Cells were then washed with PBS-T/1% FCS, blocked for 30 min at RT using 5% BSA/PBS-T/1% FCS, and stained with primary antibodies overnight at 4°C (γH2AX (1:2,000, ab81299, Abcam) and GFP (1:1,000, 11814460001, Roche)), This was followed by washing with PBS-T/1% FCS and staining using the Click-iT EdU Cell Proliferation Kit (Baseclick, Cy5). Afterwards, cells were washed with 3% BSA/PBS and incubated with fluorescently labelled secondary antibodies (goat anti mouse 488 (A11001) and goat anti rabbit 594 (A11012), both 1:1,000, Molecular Probes) for 1 h at RT. Cells were washed with PBS-T, stained with 0.4 µg/ml DAPI (Sigma Aldrich) in PBS for 5 min at RT and mounted using Vectashield antifade mounting media (Vector Laboratories). Representative images were taken using a Zeiss microscope and the Metafer4 software (MetaSystems) and subsequently processed using the ImageJ software, where brightness and contrast were adjusted for every picture if necessary.

#### Camptothecin treatment and cell cycle-specific DSB repair analysis

U2OS RAD54B-KO cells were treated with 5 nM Camptothecin (CPT) for 24 h. One hour before fixation, 10 µM of the thymidine analog EdU was added to analyze DSB repair in a cell cycle-specific manner. After fixation, DAPI and EdU intensities were scanned and measured using a Zeiss microscope and the Metafer4 software (MetaSystems). The cell cycle distribution of the scanned cell population was plotted for DNA content (measured by DAPI intensity) and EdU incorporation. EdU-negative G1- and G2-phase cells were differentiated based on their DAPI content. S phase cells incorporated EdU and were excluded from the analysis. EdU-negative G2-phase cells were analyzed for YFP and γH2AX. γH2AX foci were counted manually in 20-40 cells per condition and experiment.

---

**Figure S1. Related to Figure 1 and 2.**
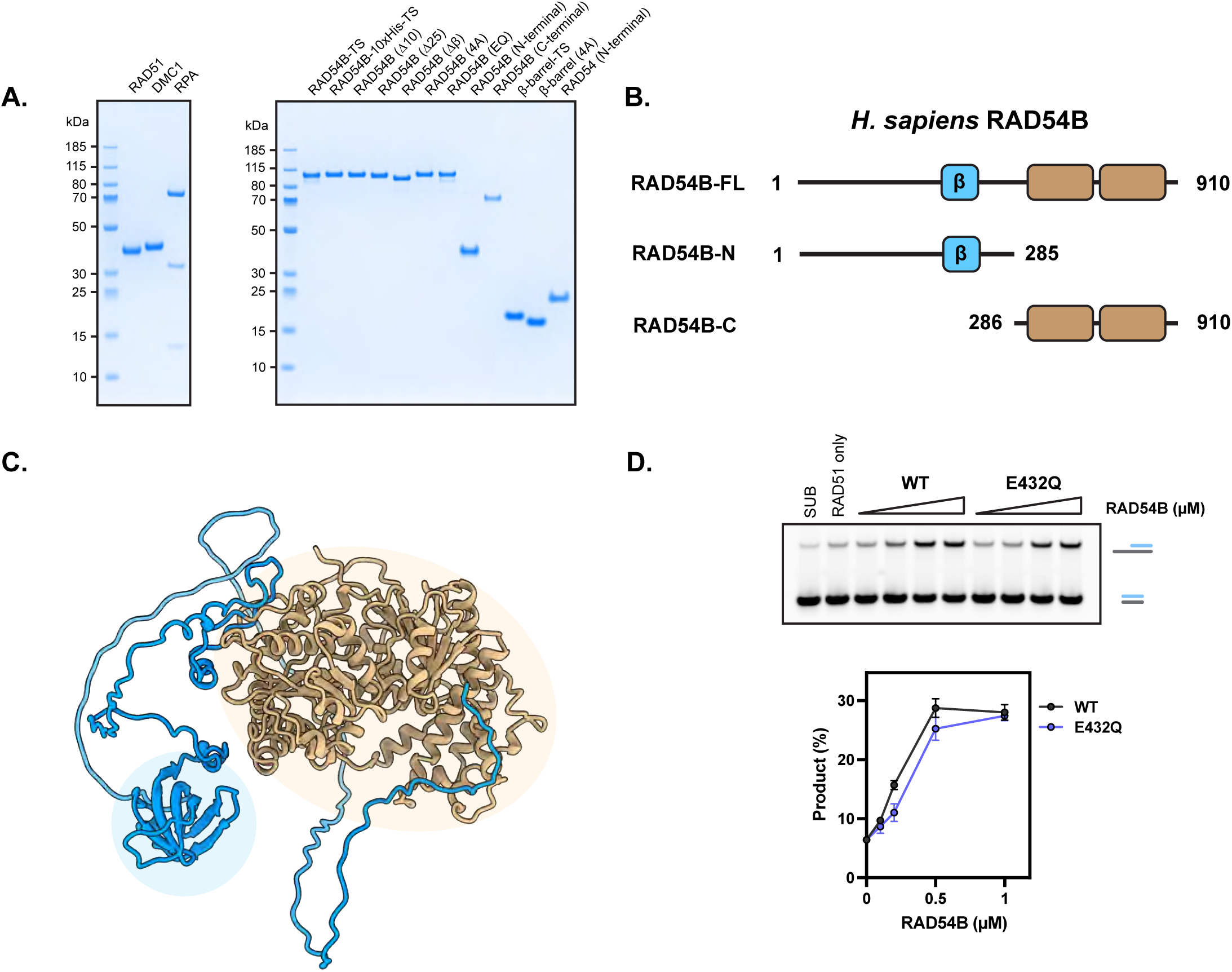
**(A)** SDS-PAGE analysis of purified proteins in this study. **(B)** Domain schematic of RAD54B. **(C)** AlphaFold3 model of human RAD54B. N-terminal region (Blue) and C-terminal region (brown). **(D)** Representative gel images showing the stimulation of RAD51-driven strand exchange by RAD54B (wild-type and mutant), averages are shown with error bars depicting standard deviation (n = 3).

**Figure S2. Related to Figure 3.**
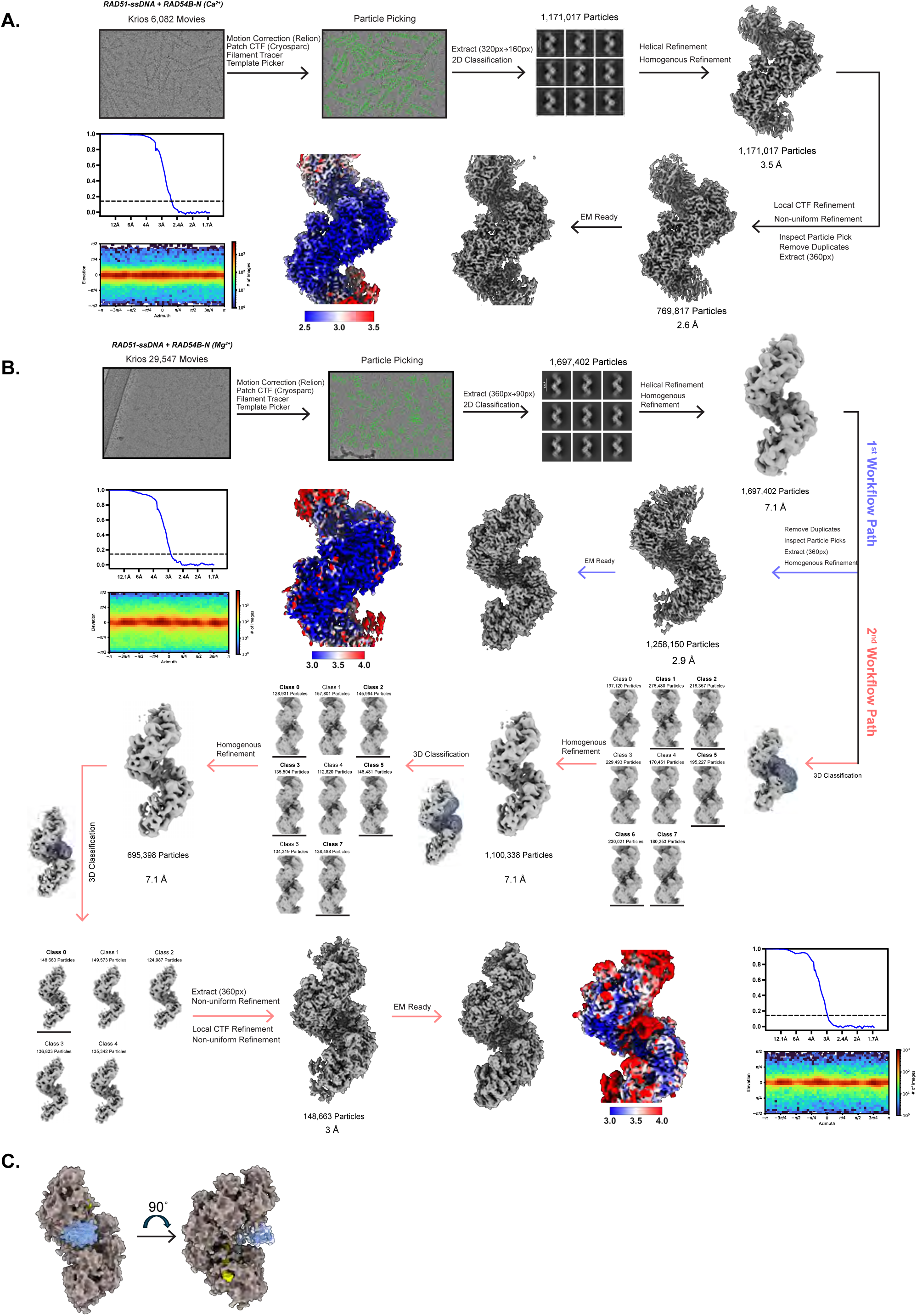
**(A)** Data processing flowchart of the Ca^2+^-ATP RAD51-ssDNA filaments bound to RAD54B-N. **(B)** Data processing flowchart of the Mg^2+^-ATP RAD51-ssDNA filaments bound to RAD54B-N. 1^st^ Workflow pathway (purple) for processing non-b-domain bound structure. 2^nd^ Workflow pathway (salmon) for processing the b-domain bound structure. **(C)** AlphaFold3 model of RAD51-ssDNA filaments with the β-domain bound.

**Figure S3. Related to Figure 3.**
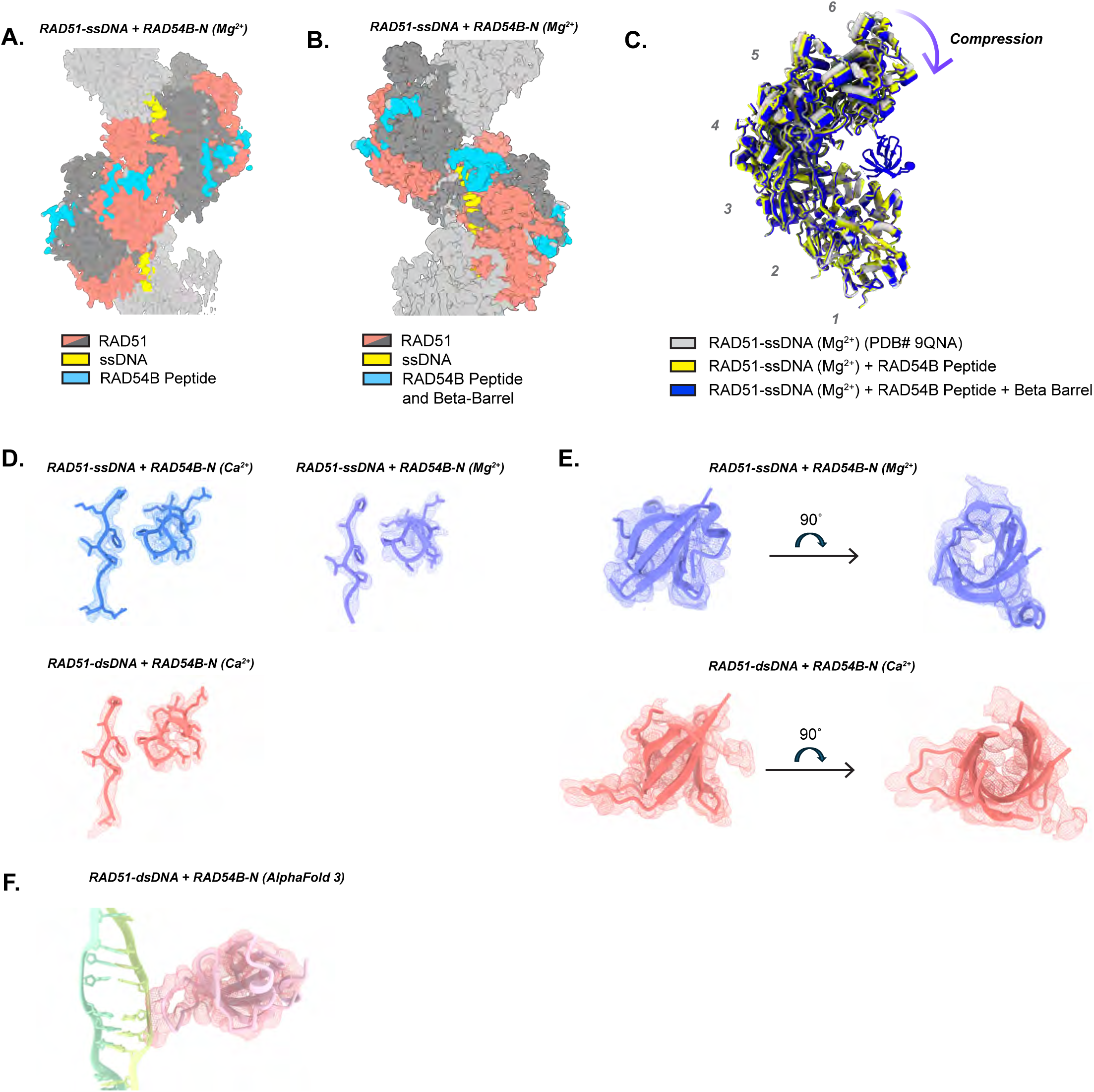
**(A)** Cryo-EM map of the Mg²⁺–ATP–bound RAD51-ssDNA filament in complex with RAD54B-N showing peptide density but no β-domain density. **(B)** Cryo-EM map of the Mg²⁺–ATP–bound RAD51–ssDNA filament in complex with RAD54B-N showing β-domain density and partial peptide density. **(C)** Structural comparison of RAD51-ssDNA filaments with or without RAD54B-N, showing filament compression upon RAD54B-N binding. Three models are aligned to the first RAD51 protomer. **(D)** Cryo-EM map quality of RAD54B sites 1 and 2 in different complexes from this study. **(E)** Cryo-EM map quality of the RAD54B β-domain in complex with RAD51-ssDNA and RAD51-dsDNA filaments. **(F)** AlphaFold3 model of RAD51-dsDNA filaments with the RAD54B β-domain (RAD51 hidden), showing how the β-domain engages dsDNA and fits the density observed in the cryo-EM map.

**Figure S4. Related to Figure 3.**
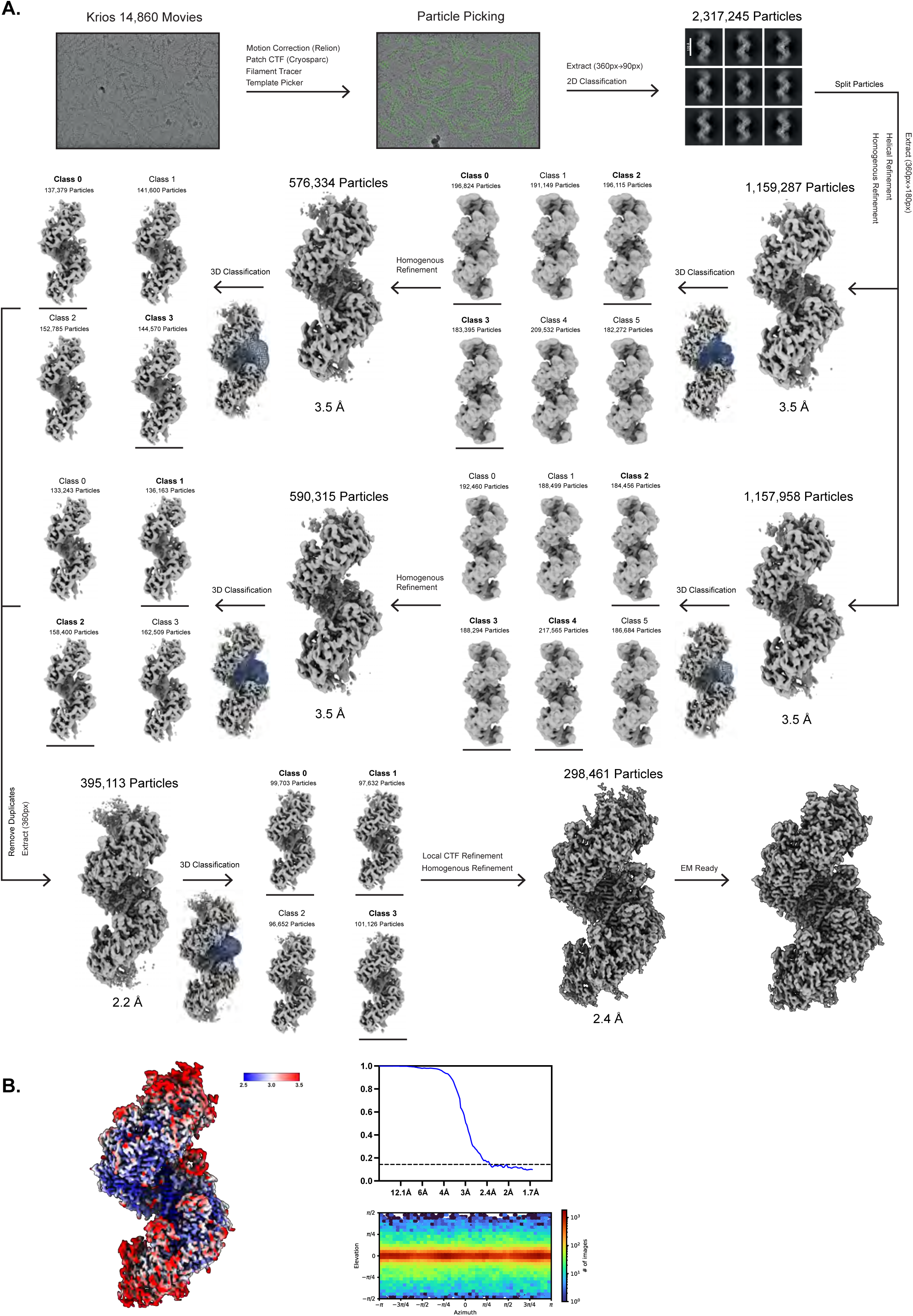
**(A)** Data processing flowchart of the Ca^2+^-ATP RAD51-dsDNA filaments bound to RAD54B-N. **(B)** Cryo-EM map of the filament from **(A)** coloured by local resolution. Orientation distribution and FSC curve of the filament from **(A)**.

**Figure S5. Related to Figures 5-6.**
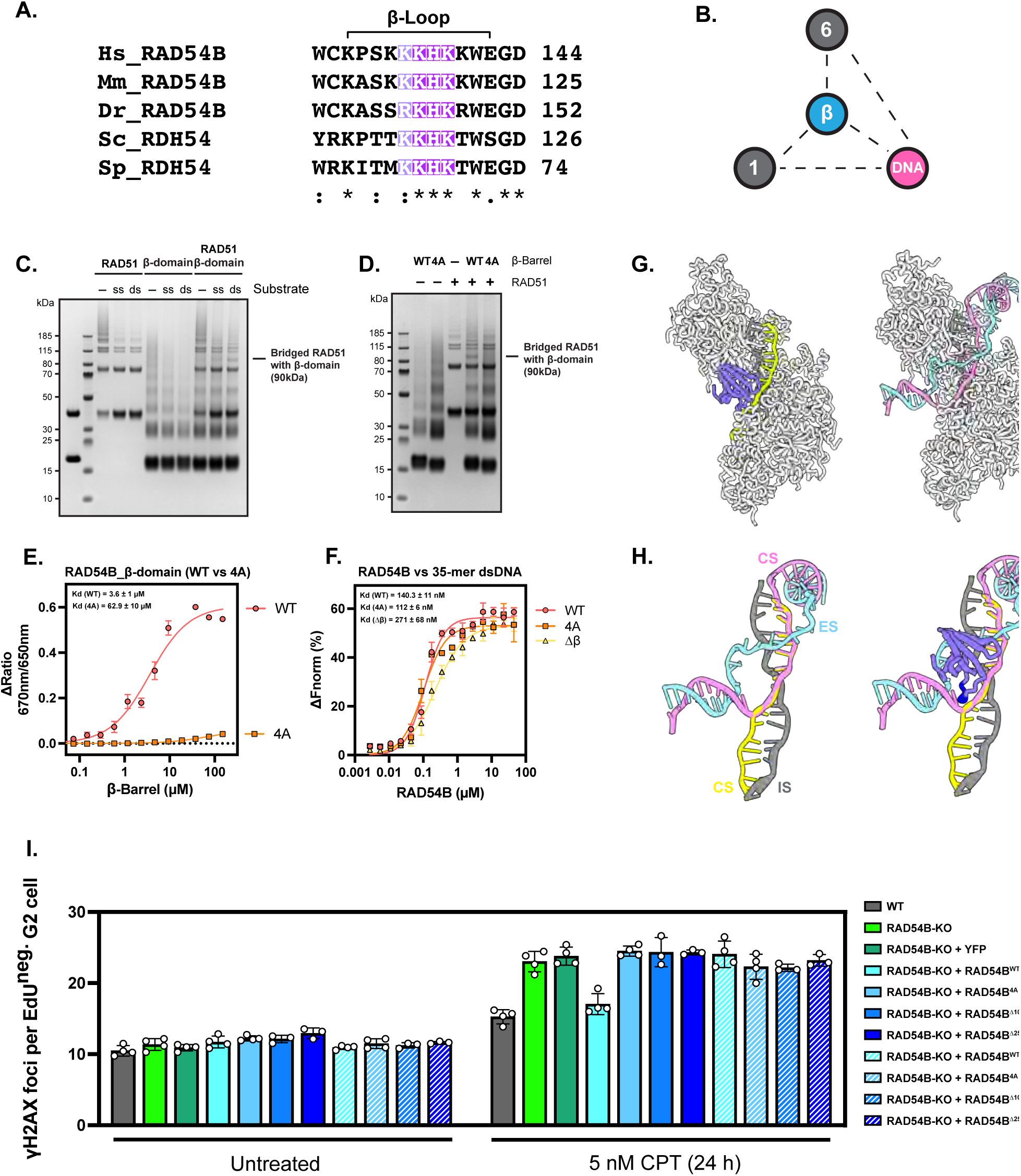
**(A)** Sequence alignment of the β-loop among RAD54B orthologs. **(B)** Schematic model of the β-domain forming a triangular interaction with two RAD51 protomers (#1 and #6) and dsDNA. **(C)** Full-size gel image corresponding to Fig. 5B. **(D)** Full-size gel image corresponding to Fig. 5D. **(E)** Microscale thermophoresis (MST) binding curves comparing wild-type RAD54B β-domain and the 4A mutant. Spectral shift curves were generated using Monolith software. **(F)** MST binding curves of full-length RAD54B (wild-type and mutants) with 35-mer dsDNA. For **(C-F)**, averages are shown with error bars depicting standard deviation (n = 3). **(G)** Structure of the RAD54B-RAD51-dsDNA filament (left) and the RAD51-D-loop complex (right; PDB: 9SW0). **(H)** Superposition of the two structures in **(G)**, aligned on RAD51, showing that the RAD54B β-domain is positioned to engage the complementary strand during strand invasion. Gray (incoming strand; IS), yellow/pink (complementary strand; CS), blue (exchanged strand; ES). **(I)** The RAD51 interaction sites and the β-domain of RAD54B are important for DSB repair by HR in U2OS cells. γH2AX foci levels in U2OS WT and RAD54B-KO cells with and without treatment with 5 nM Camptothecin (CPT) for 24 h. RAD54B-KO cells were left untransfected or were transiently transfected with constructs for YFP-only, YFP-tagged WT RAD54B, RAD54B 4A, RAD54B Δ10 or RAD54B Δ25. 1 h before fixation, 10 µM EdU was added to monitor the cell cycle. γH2AX foci were quantified in EdU-negative G2-phase cells. In transfected cells, γH2AX foci were enumerated in YFP-positive (solid bars) and YFP-negative cells (striped bars). Bars represent the mean ± SEM (n=3-4), dots represent single experiments.

**Figure S6. Related to Figure 6.**
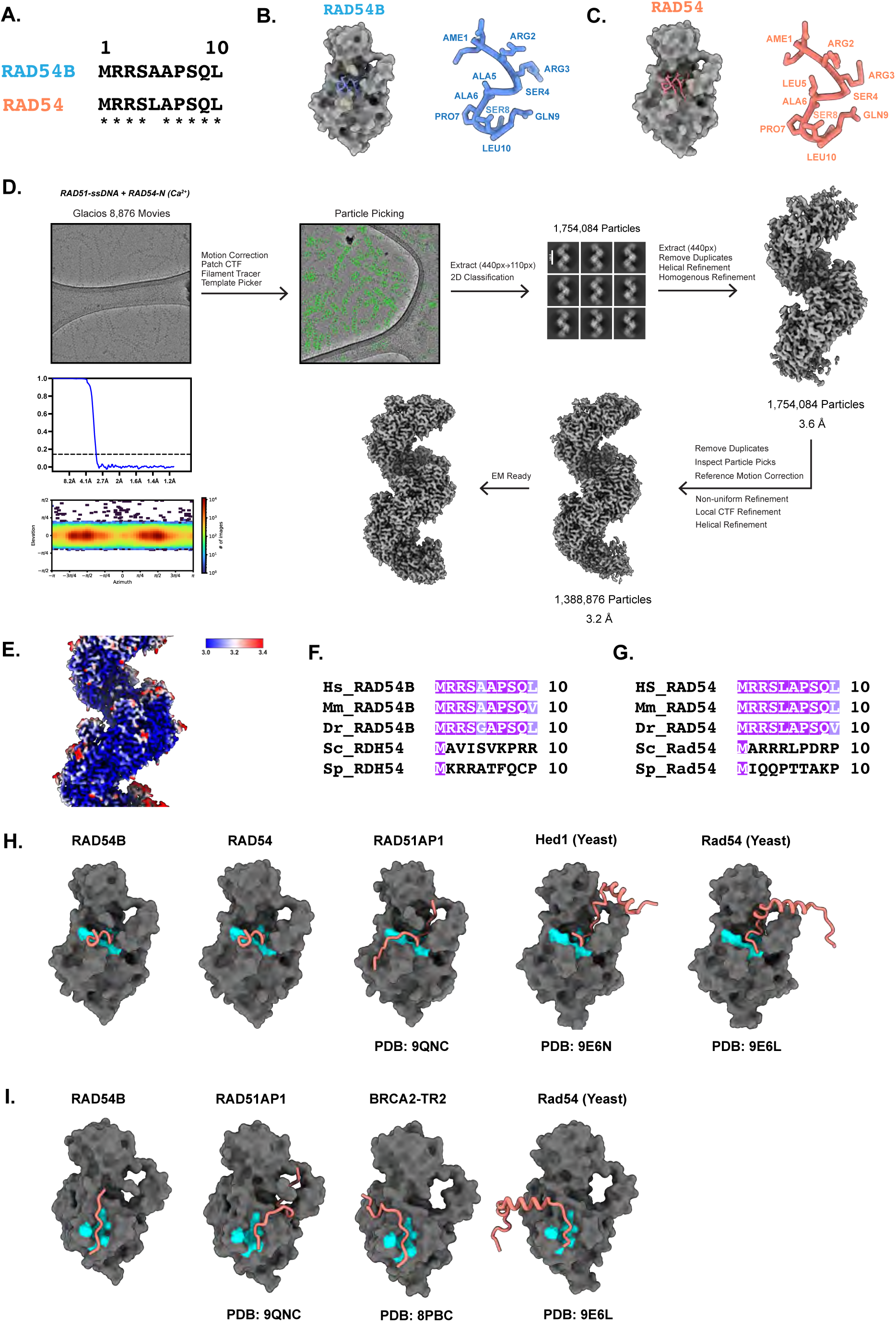
RAD54B share common RAD51 binding sites with other RAD51 modulators. **(A)** Sequence alignment of the first 10 amino acids of human RAD54 and RAD54B. **(B)** Structural model of RAD54B N-terminal residues (1–10) interacting with a RAD51 protomer. **(C)** Structural model of RAD54 N-terminal residues (1–10) interacting with a RAD51 protomer. **(D)** Data processing flowchart for the Ca²⁺–ATP RAD51-ssDNA filaments bound to RAD54-N. **(E)** Cryo-EM map of the filament from **(D)**, coloured by local resolution. **(F)** Sequence alignment of RAD54B orthologs. **(G)** Sequence alignment of RAD54 orthologs. **(H)** Comparison of RAD51 modulators known to bind the RAD51 N-terminal pocket (site 1). **(I)** Comparison of RAD51 modulators known to bind the FxPP motif–binding pocket (site 2).

## Notes

### Competing Interest Statement

The authors have declared no competing interest.

